# ZDHHC20 palmitoylates TAP1-TAP2 antigen peptide transporter to inhibit MHC-I presentation and promotes tumor immune evasion

**DOI:** 10.64898/2026.11.18.767107

**Authors:** Yizhen Jin, Junko Matsuzaki, Haochi Chen, Xuan Lu, Jianqun Liao, Feng Qian, Yekaterina Galat, Takemasa Tsuji, Jefferson Baker, Seungmin Yeom, Lei Wang, Yan-Chun Li, Kunle Odunsi, Hening Lin

**Affiliations:** Department of Medicine, The University of Chicago, 900 E. 57th Street, Chicago, IL 60637, USA; Department of Chemistry, The University of Chicago, 900 E. 57th Street, Chicago, IL 60637, USA; Graduate Program of Biochemistry, Molecular and Cell Biology, Department of Molecular Biology and Genetics, Cornell University, Ithaca, NY 14853, USA; University of Chicago Comprehensive Cancer Center, The University of Chicago, 900 E. 57th Street, Chicago, IL 60637, USA; Department of Obstetrics and Gynecology, The University of Chicago, 900 E. 57th Street, Chicago, IL 60637, USA; Howard Hughes Medical Institute; Department of Medicine and Department of Chemistry, The University of Chicago, 900 E. 57th Street, Chicago, IL 60637, USA

## Abstract

Understanding mechanisms governing the sensitivity of tumors to immunotherapy is crucial to improving cancer treatment. Protein cysteine palmitoylation plays important roles in immune regulation. Analyzing available human patient data, we identified ZDHHC20 as a key negative regulator of anti-tumor immunity, exhibiting a strong inverse correlation with CD8 T-cell infiltration. ZDHHC20 knockout markedly increased MHC-I expression. Mechanistically, we identified the antigen peptide transporter complex TAP1–TAP2 as previously unrecognized substrates of ZDHHC20. ZDHHC20–mediated palmitoylation of TAP1/2 promotes their ubiquitination by E3 ubiquitin ligases RNF5 and RNF185 and ER-associated degradation. ZDHHC20 loss in melanoma and ovarian cancer cells enhanced MHC-1 antigen presentation and CD8 T-cell activation, and suppressed tumor growth in immunocompetent mice. ZDHHC20 loss synergizes with anti–PD-1 immunotherapy, resulting in markedly enhanced inhibition of B16-F10 melanoma tumor in mice. These findings establish ZDHHC20 as a promoter of tumor immune evasion and a promising target to enhance the efficacy of immune checkpoint blockade.

**Significance:** This study establishes a previously unrecognized link between protein cysteine palmitoylation and MHC-I antigen presentation. It also identifies a new mechanism via which cysteine palmitoylation regulates ER-associated degradation (ERAD), revealing that palmitoylation can direct folded proteins for regulated ERAD. The study defines a tumor-intrinsic, post-translational mechanism of immune evasion in which DHHC20 reduces TAP1/2 abundance, suppresses MHC-I antigen presentation, and limits antitumor T-cell recognition. Its clinical relevance is supported by the association of high ZDHHC20 expression with resistance to anti-PD-1 therapy in patients and by the marked enhancement of anti-PD-1 efficacy following ZDHHC20 deletion in mouse tumor models. These findings identify DHHC20 as a potential therapeutic target for restoring tumor antigen presentation and improving responses to immune checkpoint blockade.

## Introduction

Immune checkpoint blockade (ICB) has fundamentally reshaped cancer therapy, marking an evolutionary leap in our ability to harness the immune system to treat cancer^1^. Yet only a minority of patients achieve durable clinical responses, underscoring an urgent need to identify mechanisms of resistance and develop strategies to enhance ICB efficacy^2^. Effective antitumor immunity relies on the recognition of tumor antigens presented by major histocompatibility complex class I (MHC-I) molecules by cytotoxic CD8^+^ T cells^3^. Through MHC-I antigen presentation, intracellular peptides derived from mutated proteins are processed by the proteasome, transported into the endoplasmic reticulum (ER) by the transporter associated with antigen processing (TAP) complex, and subsequently loaded onto MHC-I molecules for surface presentation^4^. This process is essential for immune surveillance that allows CD8 T cells to infiltrate, identify, and eliminate tumor cells. However, many tumors develop strategies to downregulate or disrupt components of the MHC-I pathway, leading to impaired antigen presentation and immune escape^5^. Such defects in antigen processing and presentation are strongly associated with poor responses to ICB therapies and reduced patient survival across cancer types^6,7^.

Protein S-palmitoylation, a reversible lipid modification catalyzed by the ZDHHC family of palmitoyl acyltransferases, has recently emerged as a critical post-translational mechanism regulating immune signaling^8^. Dysregulated palmitoylation has been increasingly linked to cancer progression and immune evasion^9,10^. For example, ZDHHC9 has been shown to palmitoylate PD-L1 in tumor cells, thereby stabilizing PD-L1 and suppressing CD8 T cell activity^11^. Additionally, ZDHHC9 palmitoylates TIM-3 in CD8 T cells and NK cells—preventing its polyubiquitination and degradation, which promotes immune exhaustion^12^. Likewise, ZDHHC3 has been reported to palmitoylate the immune checkpoint PD-L1 and B7-H4, blocking their lysosomal degradation and sustaining immunosuppression^13,14^.

In this study, by systematically analyzing the correlation between the levels of the 23 human *ZDHHC* genes and tumor infiltrating cytotoxic CD8 T level, we identified *ZDHHC20* as a potential anti-tumor immunity suppressor gene. We identified TAP1 and TAP2 as novel substrates of ZDHHC20 and demonstrated that ZDHHC20, by palmitoylating TAP1/2 complex, reduced the level of TAP complex. Mechanistically, palmitoylation of TAP1/2 recruits RNF5/185, which leads to increased ubiquitination and ER-associated degradation (ERAD). Consequently, ZDHHC20 suppressed MHC-I surface expression and antigen presentation, thereby diminishing CD8 T-cell activation and promoting tumor immune evasion. Genetic ablation of *ZDHHC20* increased MHC-I expression and enhanced antitumor T-cell responses both *in vitro* and *in vivo*. Together, our findings uncover a previously unrecognized ZDHHC20–TAP1/2–MHC-I axis that regulates tumor immune visibility and provide mechanistic insights into how palmitoylation contributes to immune escape, highlighting ZDHHC20 as a therapeutic target to sensitize tumors to immunotherapy.

## Results

### ZDHHC20 regulates MHC-I level in multiple cancer cell lines

We systematically ranked all S-palmitoyl transferase ZDHHC family members based on their correlation with CD8 T-cell infiltration across various tumor types^15^ (**Figure 1A**). The rationale was that if the expression of a given *ZDHHC* gene negatively correlates with CD8 T-cell infiltration across multiple cancer types, it may function as an anti-tumor immunity suppressor within the tumor microenvironment. The top five candidate immune-suppressive members identified were *ZDHHC9*, *ZDHHC20*, *ZDHHC7*, *ZDHHC3*, and *ZDHHC5*. We further ranked these candidates by averaging the negative correlation slopes across tumor types (**Figure 1B**). Notably, several previous studies provided strong experimental support for our bioinformatic observations as *ZDHHC*9/7/3/5 were recaptured as top anti-tumor immunity suppersors^11–14,16^. However, the immune suppressive function of tumor intrinsic *ZDHHC20* and its molecular mechanism remains unclear. Tomić et. al. reported that *ZDHHC20* deletion suppresses PDAC lung metastasis *in vivo* through effects on the innate immunity^17^. In another recent study, Lu et. al. reported that ZDHHC20 palmitoylates CD80 in the antigen presenting cells which help stabilize CD80 for the T-cell co-stimulation effect^18^.

**Figure 1.**
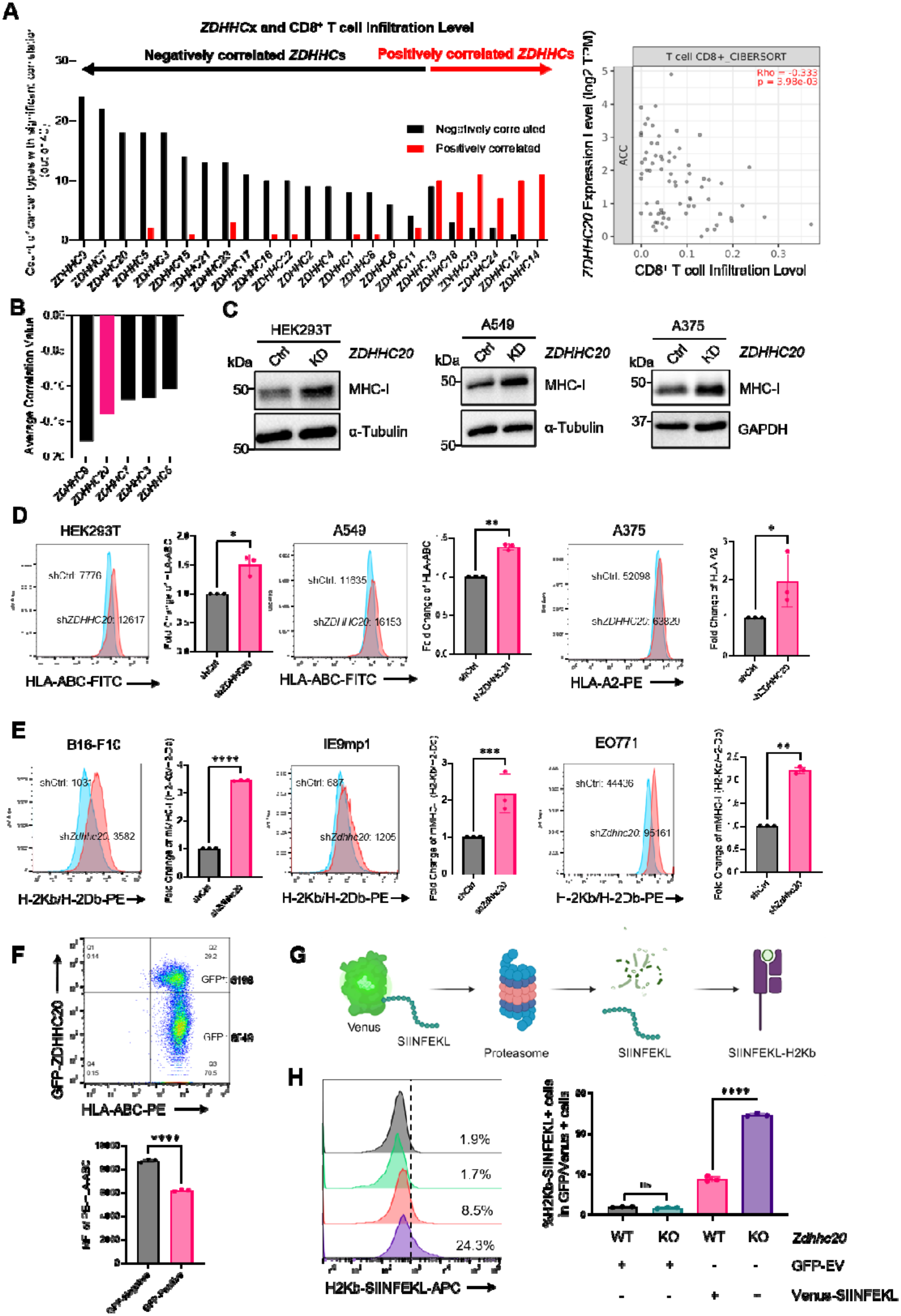
*ZDHHC20* negatively correlates with CD8 T-cell infiltration and downregulates MHC-I levels in multiple cancer cell lines. **A**. Correlation analysis between tumor-infiltrating CD8 T-cell levels and expression of each *ZDHHC* family gene across cancer types. The number of cancer type showing a negative correlation is plotted for each *ZDHHC* gene; *ZDHHC20* is highlighted in pink. An example correlation plot of *ZDHHC20* expression level and CD8^+^ T cell infiltration level is shown on the right. **B**. Average correlation coefficients (negative-slope values) for the top five *ZDHHC* genes ranked by negative correlation with CD8 T-cell infiltration. **C**. Western blot of total MHC-I levels in HEK293T, A549, and A375 cell lines following *ZDHHC20* KD. **D**. Flow cytometric analysis of surface MHC-I levels following *ZDHHC20* KD (N = 3). **E**. Flow cytometric analysis of surface MHC-I levels following *Zdhhc20* KD in mouse IE9mp1, B16-F10, and EO771 cell lines (N = 3). **F**. Flow cytometric analysis of total MHC-I levels in HEK293T cells transfected with GFP-ZDHHC20. Cells were gated into GFP-positive and GFP-negative populations, and the mean fluorescence intensity (MFI) of HLA-ABC staining was compared between the two populations. **G**. Scheme showing the SIINFEKL-based MHC-I presentation system. **H**. Representative flow cytometry plots and quantification of H2Kb-SIINFEKL percentage following *Zdhhc20* KO in B16-F10 cells. Data are presented as mean ± SD. Statistical analysis: paried t test: (D, E, F). unpaired t test: (H). Sample sizes: (D, E, H) n = 3 independent experiments. Images in C show representative results from two independent experiments.

To explore the role of ZDHHC20 in anti-tumor immunity, we analyzed the potential substrate list identified using the Bump-hole strategy for ZDHHC20^19,20^. Interestingly, several antigen presentation pathway (APP) related proteins were identified as potential ZDHHC20 substrates including HLA-C, TAP1, TAP2, TAPBP and BCAP31 (**Figure S1A**). Based on this, we hypothesized that ZDHHC20 may regulate MHC-I. To test this, we first generated stable *ZDHHC20* knock-down (KD) human cell lines in HEK293T, A549, and A375 cells (**Figure S1B**). Western blot analysis showed that *ZDHHC20* KD cells exhibited increased total MHC-I expression compared with controls (**Figure 1C**). We then measured cell surface MHC-I by assessing human HLA-ABC expression via flow cytometry (HLA-A2 for A375 cells). *ZDHHC20* KD cells displayed significantly higher levels of MHC-I on the cell surface (**Figure 1D**). To assess whether the inhibitory effect of *ZDHHC20* is conserved in mouse cancer cells, we generated stable *Zdhhc20* KD murine cancer cell lines (**Figure S1B**). Similarly, *Zdhhc20* KD resulted in a significant elevation of surface MHC-I expression in these mouse cells (**Figure 1E**). In addition, ZDHHC20 overexpression reduced cell-surface MHC-I levels in HEK293T cells (**Figure 1F**). To confirm that the increase in MHC-I was functionally meaningful, we employed a SIINFEKL-based antigen-presentation system (**Figure 1G**)^21^. Specifically, we used mouse B16-F10 cells overexpressing Venus-SIINFEKL and compared the surface levels of the H-2K -SIINFEKL complex between control and *Zdhhc20* CRISPR Knockout (KO) cells. The percentage of H-2K -SIINFEKL positive cells was significantly higher in the *Zdhhc20* KO lines (**Figure 1H**). Since MHC-I is broadly expressed across most cell types and plays a pivotal role in antigen-presenting cells, we sought to determine whether ZDHHC20 regulates MHC-I in antigen presenting cells (APC). To this end, we generated stable *Zdhhc20* KD mouse macrophage and dendritic cell lines. Similarly, *Zdhhc20* KD resulted in a significant increase in surface MHC-I levels in the APC cells (**Figure S1C**).

Cancer cells have long been known to down-regulate MHC-I through multiple mechanisms, such as suppressing the transcription of antigen-processing and presentation pathway (APP) related genes and enhancing MHC-I degradation to facilitate immune evasion^22^. We observed no significant difference in the transcriptional level of APP-related genes using qPCR between control and *ZDHHC20* KD HEK293T cells, suggesting the increased MHC-I level is not due to transcriptional change (**Figure S1D**). We also excluded the difference of MHC-I lysosomal degradation level between control and *ZDHHC20* KD (**Figure S1E**).

Notably, both mRNA and protein level of *ZDHHC20* is upregulated in most cancer types compared to the nearby normal tissues (**Figure S1F and Figure S2**)^23,24^. Given that we observed that *ZDHHC20* can downregulate MHC-I level in multiple cancer cell lines and its expression level correlates with CD8 T-cell infiltration, we hypothesized that the ZDHHC20–MHC-I axis may represent a novel mechanism of cancer immune evasion.

### ZDHHC20 palmitoylates TAP1/2 complex to regulate MHC-I

We next investigated how ZDHHC20 regulates MHC-I expression. Because multiple MHC-I-associated proteins were identified in the bump-hole substrate screen, we first evaluated each candidate individually. We excluded HLA-C, as it was not validated as a substrate of ZDHHC20 (**Figure S3**). We also confirmed that BCAP31 is a bona fide substrate of ZDHHC20 and identified Cys23 as its palmitoylation site which is consistent as previous reports (**Figure S4A**)^19^. However, re-expression of either wild-type (WT) BCAP31 or the palmitoylation-deficient C23A mutant in *BCAP31* KO HEK293T cells did not produce a significant difference in MHC-I expression (**Figure S4B**). These findings suggest that BCAP31 palmitoylation is unlikely to account for the effect of ZDHHC20 on MHC-I regulation. We therefore focused our subsequent studies on the TAP complex.

TAP complex, a member of the ATP-binding cassette (ABC) transporter family, plays a crucial role in the processing and presentation of MHC-I restricted antigens^25^. Given this distinctive role, we investigated whether the TAP1/2 transporter are substrates for ZDHHC20. The TAP complex comprises two subunits: TAP1 and TAP2, both containing 10 transmembrane regions across the ER membrane^25–30^. We first confirmed that both TAP1 and TAP2 are palmitoylated by ZDHHC20 but not by the catalytic dead C156S mutant (ZDHHS20) using an Alk14 labeling assay (**Figure 2A**). Additionally, acyl-biotin exchange (ABE) assay verified that endogenous TAP1 and TAP2 undergo ZDHHC20-mediated palmitoylation (**Figure 2B**).

**Figure 2.**
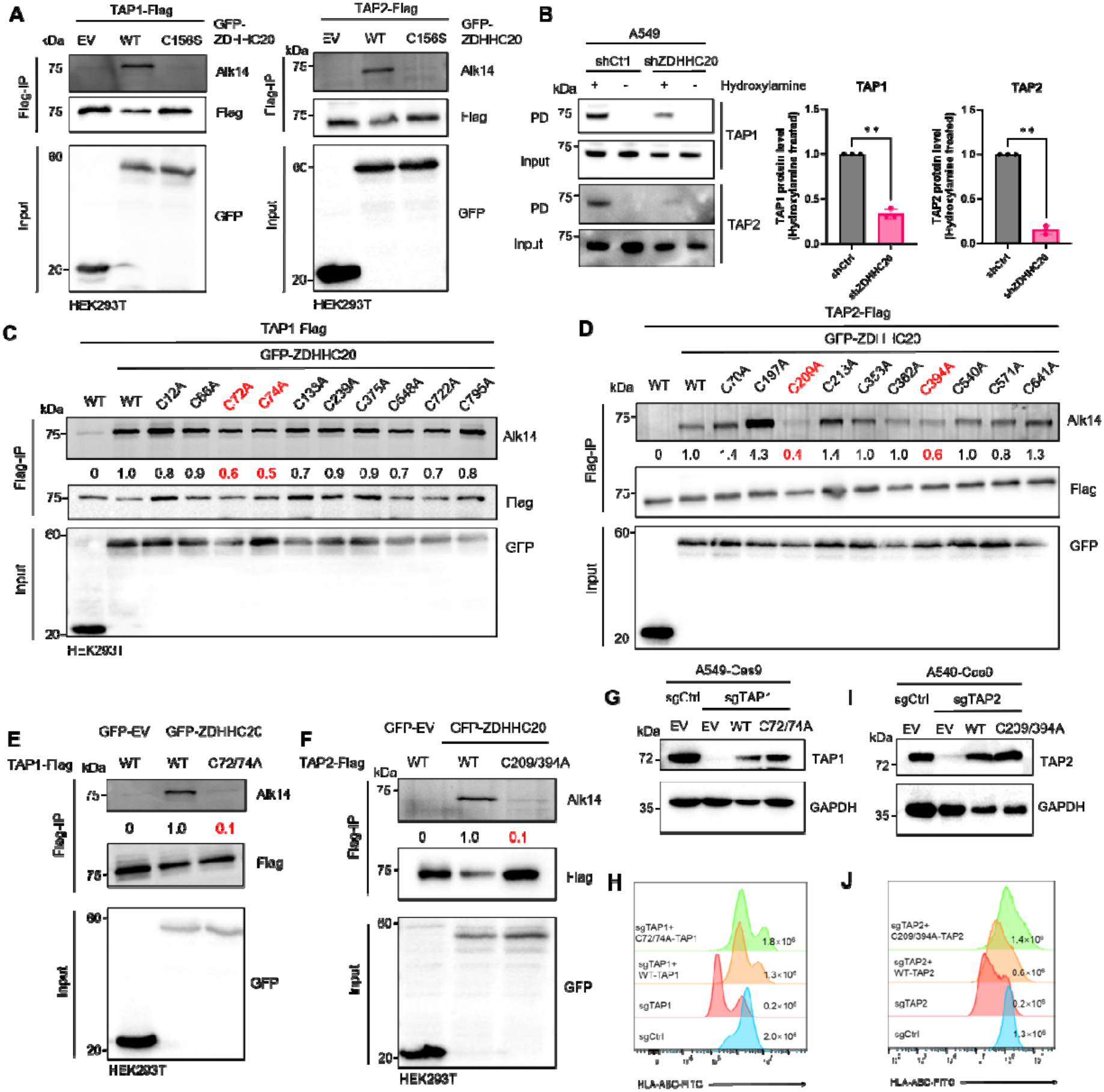
ZDHHC20 palmitoylates TAP1/TAP2 complex to regulate MHC-I level. **A**. Alk14 labeling assay of TAP1 and TAP2 co-overexpressed with EV, WT-DHHC20 and catalytic dead C156S mutant (DHHS20). **B**. ABE assay in WT and ZDHHC20 KD A549 cells and quantification of TAP1/2 protein level. **C-D**. Alk14 labeling assay of TAP1(**C**) and TAP2(**D**) Cysteine to Alanine mutants co-overexpressed with WT ZDHHC20 for palmitoylation site identification. Palmitoylation sites are highlighted in red. **E-F**. Alk14 labeling assay of WT and mutant TAP1 (E)/TAP2(F) co-overexpressed with WT ZDHHC20. **G**. Re-introduction of WT or C72/74A-TAP1 back into TAP1 KO A549 cells. **H**. C72/74A-TAP1 restored higher level of surface MHC-I level than WT-TAP1. **I.** Re-introduction of WT or C209/394A-TAP2 back into TAP2 KO A549 cells. **J**. C209/394A-TAP2 restored higher level of surface MHC-I level than WT-TAP2. **K**. AlphaFold3 prediction of TAP1/2 complex with palmitoylation. Palmitoyl groups are highlighted in green. Image in A, B, E, H, J shows representative results from at least two independent experiments.

Both TAP1 and TAP2 possess ten cysteine residues located either in the transmembrane domains or cytosolic regions, making them potential sites for palmitoylation. To identify the specific cysteine residues modified by ZDHHC20, we performed site-directed mutagenesis, substituting each cysteine with alanine, and compared the Alk14 labeling levels to those of the WT TAP1/2. For TAP1, mutations at Cys72 and Cys74 (C72A and C74A) resulted in 40-50% decrease in Alk14 labeling signal (**Figure 2C**). For TAP2, mutations at Cys209 and Cys394 (C209A and C394A) led to 60% and 50% reduction in Alk14 labeling signal, respectively (**Figure 2D**). We then generated double mutants for both TAP1 (TAP1_C72/74A) and TAP2 (TAP2_C209/394A), which further diminished Alk14 labeling signals (**Figure 2E**). Our results indicate that ZDHHC20 catalyzes the palmitoylation of TAP1 primarily at Cys72 and Cys74, and TAP2 at Cys209 and Cys394. The TAP1 construct used in this study corresponds to human TAP1 isoform 2 (UniProt Q03518-2, 808 amino acids), rather than the canonical isoform 1 (UniProt Q03518-1, 748 amino acids). Isoform 2 contains a 60-residue N-terminal extension with an additional cysteine, bringing its total cysteine count to 10^31^. We selected this isoform to include the additional potential palmitoylation site absent from isoform 1 (**Figure S5A**). To determine whether ZDHHC20 also palmitoylates canonical TAP1 at the corresponding cysteine residues, we performed Alk14 labeling of wild-type isoform 1 and its C12/14A mutant. The C12/14A mutation markedly reduced ZDHHC20-dependent Alk14 incorporation, supporting Cys12 and Cys14 (corresponding to Cys72 and Cys74 in isoform 2) as the major sites of ZDHHC20-mediated palmitoylation in isoform 1 (**Figure S5B**).

Given the inhibitory role of ZDHHC20 on MHC-I level, we hypothesized that ZDHHC20-mediated palmitoylation also inhibits the TAP complex. To test this, we generated *TAP1* CRISPR-KO and *TAP2* CRISPR-KO HEK293T cells (Cas9 background, **Figure S6A and S6C**). Both *TAP1* and *TAP2* KO significantly downregulated MHC-I level as previously reported (**Figure S6B and S6D**)^32^. We then transiently knocked down *ZDHHC20* and compared the MHC-I level in control and *TAP2* KO HEK293T cells (**Figure S6E**). Consistently, transient *ZDHHC20* KD increased MHC-I level in the control HEK293T cells. In contrast, *TAP2* KO blocked this effect, which indicates the effect of ZDHHC20 on MHC-I is through the TAP complex (**Figure S6F**).

Since loss of *ZDHHC20* increases MHC-I level, the palmitoylation-deficient mutant of TAP1 and TAP2 should phenocopy loss of *ZDHHC20* and increase MHC-I levels if our hypothesis is correct. To test this, we utilized A549 cell, a lung pulmonary adenocarcinoma cell line, which express higher levels of *ZDHHC20* than HEK293T, to better demonstrate the difference between WT and palmitoylation-deficient mutants. To this end, we generated *TAP1* and *TAP2* CRISPR KO A549 cells and re-introduced either WT or palmitoylation-deficient mutant TAP1 or TAP2 into the corresponding *TAP1* or *TAP2* KO A549 cells, and then compared the MHC-I surface levels (**Figure 2G and 2I**). As expected, the palmitoylation deficient mutant of TAP1 or TAP2 restored higher MHC-I level than the corresponding WT constructs (**Figure 2H and 2J**).

Interestingly, mouse Tap1 lacks the extended N-terminal region present in human TAP1, where the palmitoylated Cys72 and Cys74 are located. In contrast, the palmitoylated cysteines at positions 209 and 394 are conserved in the mouse Tap2 sequence (**Figure S6G–H**). This difference suggests that TAP2 palmitoylation may represent a more evolutionarily conserved mechanism by which ZDHHC20 regulates MHC-I.

### Palmitoylation of TAP1/2 complex recruits RNF5/185 for ubiquitination and proteasomal degradation

After confirming that ZDHHC20-mediated palmitoylation of TAP1 and TAP2 inhibits MHC-I presentation, we then sought to investigate how TAP1/2 palmitoylation affects MHC-I. Palmitoylation is known to affect protein-protein interactions. However, none of the palmitoyl groups are closely located in the heterodimer interface based on reported TAP complex structure^28,29,33^. Consistently, Co-IP assays showed that neither ZDHHC20 overexpression nor expression of palmitoylation-deficient TAP1 or TAP2 mutants altered the TAP1–TAP2 interaction (**Figure S7A**).

Interesting, in the Alk14 labeling assays, we observed that co-overexpression of WT ZDHHC20 markedly reduced the protein levels of both overexpressed TAP1 and TAP2, whereas co-overexpression of the catalytically inactive mutant ZDHHS20 had no such effect (**Figure 2A**). This finding suggests that ZDHHC20-mediated palmitoylation may regulate the stability of TAP1 and TAP2. To further investigate this, we compared endogenous TAP1 and TAP2 protein levels between control and *ZDHHC20* KD cells. In both A549 and A375 cell lines, *ZDHHC20* KD significantly increased the protein abundance of TAP1 and TAP2 (**Figure 3A**).

**Figure 3.**
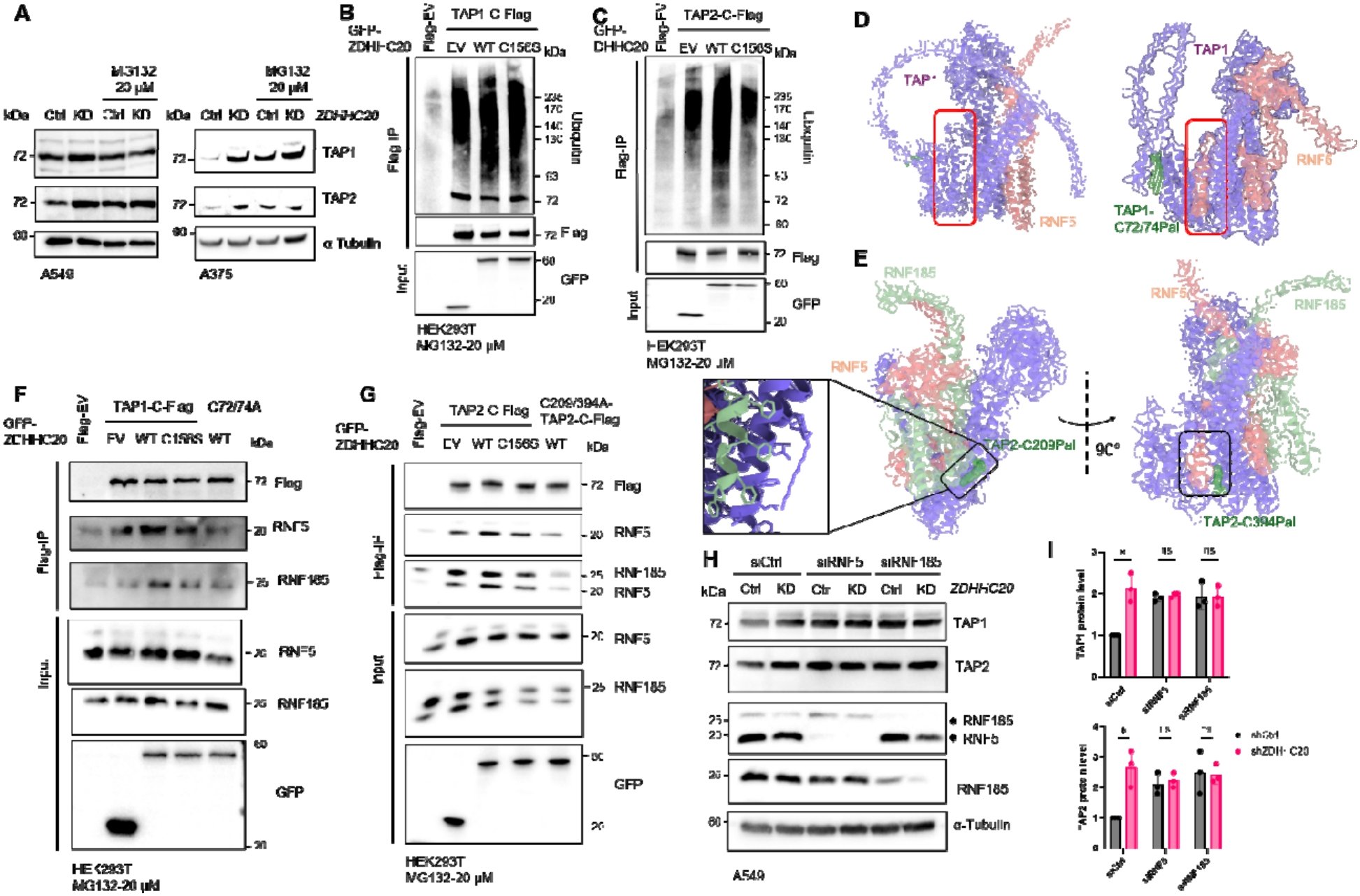
Palmitoylation of TAP1/2 complex recruits RNF5/185 for ubiquitination and leads to ERAD. **A-B**. WB analysis of TAP1 and TAP2 protein level in shCtrl and sh*ZDHHC20* A549 cells (**A**) and A375 cells (**B**) treated with or without Mg132 (20μM, 6h). C. Ubiquitination level of TAP1 and TAP2 co-overexpressed with empty vector, WT-ZDHHC20 and C156S-ZDHHC20 treated with Mg132 (20μM, 6h). **D**. AlphaFold 3 prediction of TAP1 (with or without C72/74 palmitoylation) binding pos with RNF5. **E**. AlphaFold3 prediction of TAP2 (with C209/394 palmitoylation) binding pose with RNF5/RNF185. The hydrophobic interaction interface is highlighted in black box. Surface is colored with hydrophobicity in the Zoom-in view using ChimeraX. **F-G**. Palmitoylation of TAP1 (**F**) /TAP2 (**G**) mediated by ZDHHC20 increased their interaction of RNF5/RNF185. **H**. Transient KD of RNF5 or RNF185 in shCtrl and sh*ZDHHC20* A549 cells rescued the protein level difference of TAP1 and TAP2. **I**. Quantification for TAP1/2 level for RNF5 or RNF185 KD in shCtrl and sh*ZDHHC20* A549 cells (N=3). Statistical analysis: unpaired t test: (I). Sample sizes: (I) n = 3 independent experiments. Image in (A, B, C, F, G, H) shows representative results from at least two independent experiments.

We next investigated the mechanism via which ZDHHC20-mediated palmitoylation affects TAP1 and TAP2 protein levels. Based on transcriptomic profiling of APP genes in shCtrl and sh*ZDHHC20* cells, we excluded transcriptional regulation as the cause of the TAP1/2 protein-level differences (**Figure S1E**). Since TAP1 and TAP2 have been reported to undergo proteasomal degradation^34^, we therefore examined the effect of the proteasome inhibitor MG132. Upon MG132 treatment, the differences in TAP1 and TAP2 levels between shCtrl and sh*ZDHHC20* cells were abolished, indicating that ZDHHC20-mediated palmitoylation promotes their proteasomal degradation. Consistently, co-overexpression of WT-ZDHHC20 increased the ubiquitination of both TAP1 and TAP2, whereas the ZDHHS20 mutant had no effect (**Figure 3B and 3C**).

A previous study reported that a key component of the MHC-I peptide loading complex, Tapasin (also known as TAPBP), is regulated through ERAD^35^. Specifically, the RNF185/Membralin ERAD complex recognizes unassembled TAPBP and restricts its incorporation into the peptide-loading complex (PLC), thereby modulating MHC-I surface expression^35^. We performed a similar experiment to check the ubiquitination level TAPBP with overexpression of WT-ZDHHC20. While the result suggested that the ubiquitination level of TAPBP is not regulated by ZDHHC20 (**Figure S7B**), it prompted us to hypothesize that TAP1 and TAP2 may also be subject to ERAD regulation, and that palmitoylation of TAP1/2 might act as a hydrophobic signal mimicking a misfolded region to recruit specific ERAD-associated E3 ligases for degradation^36^.

To explore whether TAP1/2 is regulated by ERAD, we curated a list of 15 E3 ligases previously implicated in the ERAD pathway^37^ and performed AlphaFold3.0^38^ based binding analysis to find the E3 ligase regulating TAP1/2. Our rationale was twofold: (A) to determine whether palmitoylation alters the binding mode between TAP1/2 and E3 ligases compared with their non-palmitoylated counterparts; and (B) to assess whether palmitoylation sites are located near the TAP–E3 interaction interfaces. In addition to the virtual interaction analysis, we cross-referenced experimentally reported TAP1 and TAP2 interactors from the BioPlex^39^ and Biogrid^40^ databases. Among the candidates, RNF5/185 emerged as potential E3 ligases for both TAP1 and TAP2. RNF5 and RNF185 are structurally homologs. Previous study reported they work together to target misfolded CFTR to ERAD^41^.

Compared with non-palmitoylated TAP1, the C72Pal/C74Pal-TAP1 structure is predicted to possess an expanded pocket between transmembrane domain (TMD) 4 and TMD8 that interacts with the C-terminal hydrophobic segment of RNF5 (residues 156–180), as highlighted in the red box in **Figure 3D**. This interaction is not observed in non-palmitoylated TAP1. In contrast, the interaction poses of WT-TAP2 and RNF5/185, as well as those between C209Pal/C394Pal-TAP2 and RNF5/185, are largely similar. However, the C-terminal phenylalanine-rich region of RNF5 (I<u>FFFF</u>WLLSI, residues 170–180) is positioned adjacent to the C394 palmitoylation site within the TAP2 TMD (**Figure 3E**), and the phenylalanine-enriched α-helical segment of RNF185 (<u>F</u>P<u>F</u>GI<u>F</u>ATA<u>F</u>, residues 142–151) lies proximal to the C209 palmitoylation group (**Figure 3E**). A zoom-in view of this dimer interface reveals four parallel phenylalanine residues aligned with the palmitoyl group, suggesting a strong hydrophobic interaction (**Figure 3E**). This hydrophobic patch may serve as a palmitoylation recognition motif for ubiquitination substrate selection.

To further test the hypothesis that TAP1/2 palmitoylation by ZDHHC20 serve as a recognition motif to recruit the ERAD E3 enzymes RNF5/185, we performed Co-IP assays to examine the interactions between TAP1/2 and RNF5/185 in the presence or absence of ZDHHC20 overexpression (**Figure 3F-G**). Notably, co-expression of WT-ZDHHC20 with either WT-TAP1 or WT-TAP2 enhanced their association with RNF5 and RNF185. In contrast, co-expression of inactive ZDHHS20 with WT-TAP1/2, or of WT-ZDHHC20 with palmitoylation-deficient TAP1/2 mutants, did not significantly alter interaction with RNF5/185 (**Figure 3F-G**). These results are consistent with the hypothesis that palmitoylation of TAP1/2 promotes recruitment of the E3 ligases RNF5 and RNF185 for ERAD. To further validate this, we transiently knocked down *RNF5* or *RNF185* in shCtrl and sh*ZDHHC20* A549 cells. As expected, knockdown of either *RNF5* or *RNF185* rescued the difference of TAP1/2 protein levels in shCtrl and sh*ZDHHC20* A549 cells (**Figure 3H-I**).

We further tested the involvement of the Phe-rich motif in RNF185 in binding to TAP2. We generated an RNF185_142–151-FA mutant (<u>A</u>P<u>A</u>G<u>IA</u>ATA<u>A</u>), in which the four phenylalanine residues within this region were substituted with alanine. We co-overexpressed WT or palmitoylation-deficient TAP2 together with either WT or the FA mutant of RNF185 in HEK293T cells. TAP2 was immunoprecipitated to assess its interaction with RNF185 and the resulting ubiquitination of TAP2 (**Figure S7C**). Consistent with our hypothesis, the palmitoylation-deficient TAP2 mutant (C209/394A) exhibited markedly reduced interaction with RNF185 compared with WT TAP2. Likewise, the RNF185 FA mutant displayed substantially diminished binding to TAP2. Correspondingly, the level of TAP2 ubiquitination closely mirrored the strength of the TAP2–RNF185 interaction for WT-TAP2. While overexpressing RNF185 clearly increased the ubiquitination of the WT TAP2, it did not obviously increase the ubiquitination of palmitoylation deficient mutant of TAP2, **Figure S7C**). In addition, we further performed the Co-IP analysis of TAP2 and RNF185 (WT or 142–151-FA mutant) with or without ZDHHC20 overexpression (**Figure S7D**). Palmitoylation of TAP2 increased its interaction with RNF185-WT but not the RNF185_142–151-FA mutant. These results further supported that the phenylalanine enriched domain is critical for recognizing palmitoylated substrates and promoting ubiquitination.

### ZDHHC20 inhibits T cell activation and effector function *in vitro*

MHC-I antigen presentation machinery is necessary for CD8 ^+^ T cell activation and function^22^. We therefore hypothesized that loss of *ZDHHC20* should promote antigen-specific CD8 T-cell responses through increased MHC-I presentation level. To test this, we co-cultured the shCtrl or sh*ZDHHC20* human melanoma cell line A375 (expressing NY-ESO1 antigen) with human peripheral T cells expressing TCR that specifically recognizes the NY-ESO-1 SLLMWITQC peptide-HLA-A*02:01 complex (**Figure 4A**)^42^. *ZDHHC20* KD in melanoma cells enhanced T-cell activation, as indicated by the increased proportion of IFN-γ CD8 T cells at the early time point and elevated levels of secreted IFN-γ after 24 hours (**Figure 4B-C**). No significant increase was observed when A375 cells were co-cultured with T cells not expressing the specific TCR, suggesting that the effect of *ZDHHC20* depends on the MHC-I–TCR axis. Concordantly, *ZDHHC20* KD A375 cells were more susceptible to T cell-mediated killing as indicated by the significant increase of early apoptosis population co-cultured with the TCR transduced T cells, while no significant difference was observed when co-cultured with untransduced T cell or without T cell (**Figure 4D-F**).

**Figure 4.**
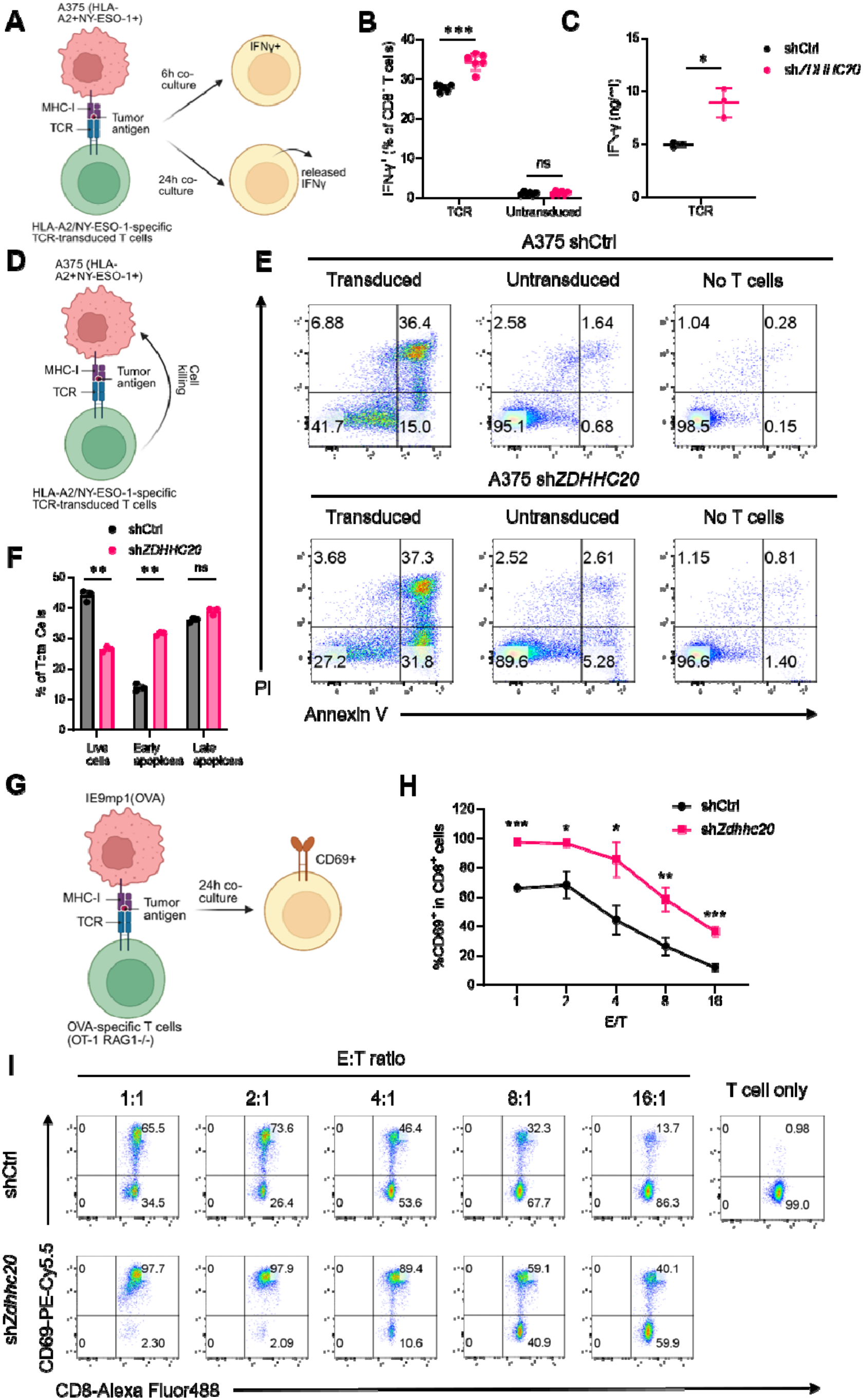
Loss of *ZDHHC20* enhances T cell activation and effector function *in vitro*. **A**. Scheme showing the human T cell co-culture assay. **B**. Flow cytometric analysis showing the percentages of IFN-γ ^+^ CD8 ^+^ T cells co-cultured with NY-ESO-1-expressing shCtrl and sh*ZDHHC20* A375 cells. **C**. ELISA quantification of IFN-γ released in the co-culture supernatants. **D**. Scheme showing the human T cell killing assay. **E**. Representative flow-cytometry plots from the killing assay. **F**. Quantification of A375 cell viability after co-culture with NY-ESO-1 TCR-T cells. Live cells were defined as PI /Annexin V, early apoptotic cells as PI /Annexin V, and late apoptotic cells as PI /Annexin V . **G**. Scheme showing the mouse T cell co-culture assay. **H**. Quantification of the CD69^+^ % in CD8^+^ T cells from the mouse T cell co-culture assay. **I**. Representative flow-cytometry plots from the mouse T cell co-culture assay. Statistical analysis: unpaired t test: (B, C, F, H). Sample sizes: (B, C, F, H) n ≥ 3 independent experiments. Images in E, I show representative results from three independent experiments.

In addition to human melanoma A375 cells, we tested whether *Zdhhc20* loss could enhance CD8 T-cell activation using OVA-expressing mouse epithelial ovarian cancer line, IE9mp1 (**Figure 4G**)^43^. Consistently, *Zdhhc20* KD in IE9mp1 cells significantly increased surface MHC-I level (**Figure 1E middle panel**). By co-culturing the H-2Kb:OVA–specific OT-1 CD8^+^ mouse T cells with IE9mp1, we confirmed that *Zdhhc20* KD significantly enhanced T cell activation under different effector cell to tumor cell ratios (E:T ratio) quantified by the CD69^+^ population (**Figure 4H-I**). Taken together, ZDHHC20 is a suppressor of cytotoxic CD8 ^+^ T cell activation through regulating MHC-I presentation level.

### Loss of *ZDHHC20* inhibits tumor growth *in vivo* through increased CD8^+^ T cell recruitment

To explore the translational relevance, we further examined the correlation of *ZDHHC20* expression and CD8^+^ T cell infiltration in different human tumor types^44^. Out of 40 tumor types, 18 showed a significant negative correlation between *ZDHHC20* expression and CD8^+^ T cell infiltration (**Figure S8A**). Moreover, correlation analysis of T cell-related genes suggested a frequent negative correlation between *ZDHHC20* expression and *CCL5* and *CD3D* expression in multiple tumor types (**Figure S8B-C**). Consistent with the co-culture experiment *in vitro*, melanoma cancer is indeed identified as a promising cancer type in which CD8^+^ T cell infiltration is significantly negatively correlated to *ZDHHC20* level in tumor. Therefore, we sought to examine the role of *Zdhhc20* for tumor growth *in vivo* using the B16-F10 murine melanoma model (**Figure 5A**).

**Figure 5.**
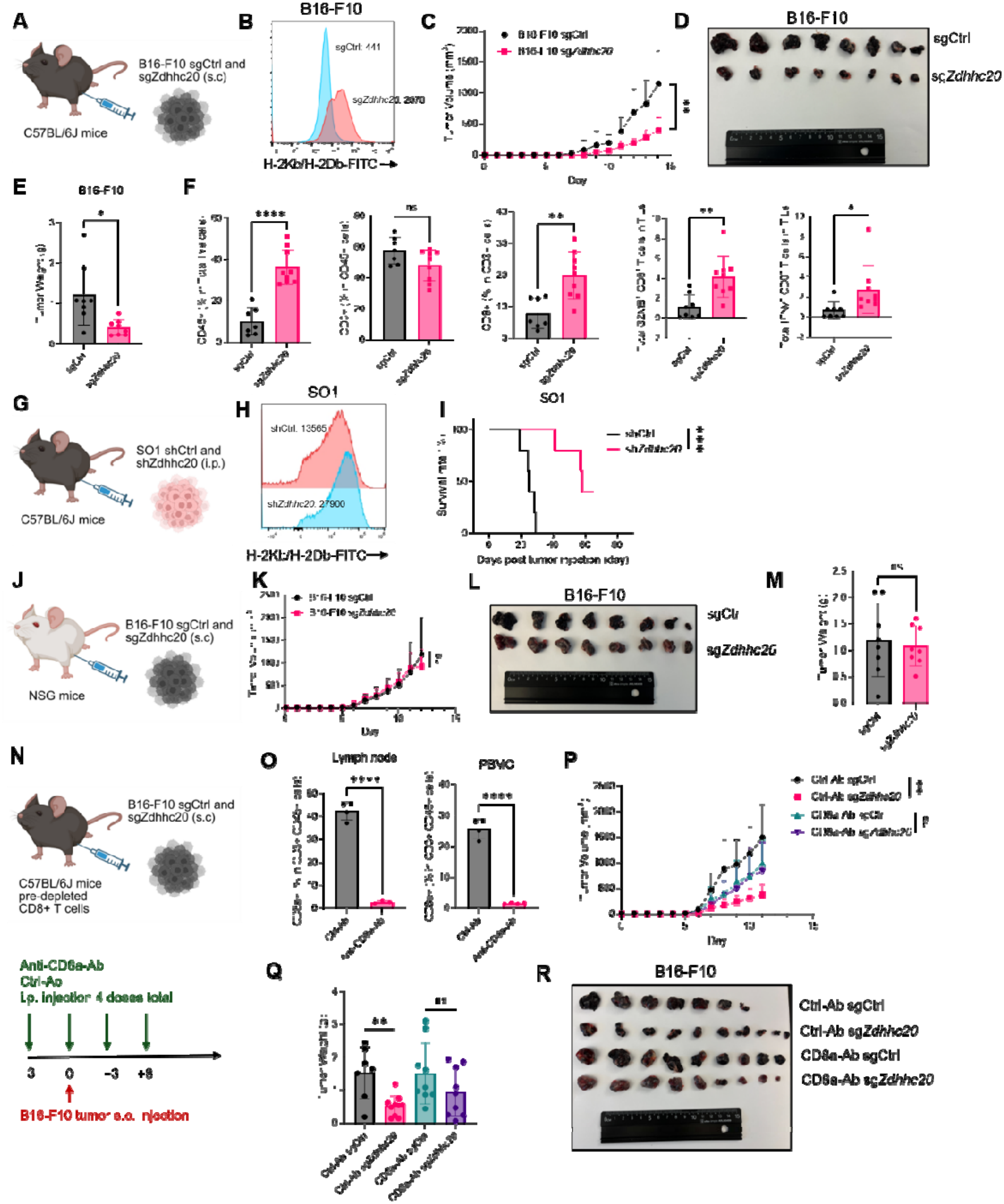
Loss of *Zdhhc20* inhibits cancer growth in vivo through increased CD8^+^ T cell recruitment. **A**. B16-F10 subcutaneous tumor model using sgCtrl and sg*Zdhhc20* B16-F10 cells in C57BL/6J mice (N = 8 for each group). **B**. Representative flow cytometry results of surface MHC-I level in sgCtrl and sg*Zdhhc20* B16-F10 cells. **C-E**. Tumor growth curve (**C**), tumor images (**D**), and tumor weight (**E**) of sgCtrl and sg*Zdhhc20* B16-F10 subcutaneous tumors in C57BL/6J mice. **F**. Flow cytometric analysis of TILs in sgCtrl and sg*Zdhhc20* B16-F10 subcutaneous tumors in C57BL/6J mice. **G**. SO1 intraperitoneal ovarian tumor model using shCtrl and sh*Zdhhc20* SO1 cells (N = 5 for each group). **H**. Representative flow cytometry results of surface MHC-I level in shCtrl and sh*Zdhhc20* SO1 cells. **I**. survival curve of SO1 shCtrl and sh*Zdhhc20* ovarian tumor model. **J-M**. B16-F10 subcutaneous tumor model using sgCtrl and sg*Zdhhc20* B16-F10 cells in NSG background mice (N = 8 for each group). Tumor growth curve (**K**), tumor images (**L**), and tumor weight (**M**) are shown. **N-T.** B16-F10 subcutaneous tumor model using sgCtrl and sg*Zdhhc20* B16-F10 cells in C57BL/6J mice with CD8^+^ T cell depletion (N = 7 for Ctrl-Ab sgCtrl, N = 10 for Ctrl-Ab sg*Zdhhc20*, N = 9 for CD8-Ab sgCtrl/sg*Zdhhc20*). The discrepancy in sample size of Ctrl-Ab treated groups resulted from the exclusion of three tumors specifically from caliper-based tumor-growth analysis and final tumor weight measurement because intraperitoneal tumor growth prevented reliable measurement. Anti-CD8a and isotype control antibodies were injected through i.p. for four doses in total at Day -3, 0, 3 and 6. Flow cytometric analysis of total CD8^+^ T cell amount in PBMC and tumor drained lymph nodes of mice receiving Ctrl-Ab and Anti-CD8a-Ab at D12 (**O**). Tumor growth curve (**P**), tumor weight (**Q**), and tumor images (**R**) are shown. Statistical analysis: unpaired t test: (C, E, F, K, M, O, P, Q, R, S); logrank (Mantel-Cox) test: (I).

We first confirmed that there is no growth difference between control or *Zdhhc20* KO B16-F10 cells when cultured *in vitro* for 72 hours with three different starting cell numbers (**Figure S8D**). Furthermore, using the DepMap database, we confirmed that CRISPR KO of *ZDHHC20* did not produce a significant effect across diverse cancer lineages (**Figure S8E**)^45^. *Zdhhc20* KO B16-F10 cells present ∼4-fold higher total MHC-I than the control group (**Figure 5B**). We then injected 0.1 million B16-F10 melanoma cells subcutaneously into C57BL/6J mice and compared the growth curve of control and *Zdhhc20* KO tumors. As expected, loss of *Zdhhc20* in B16-F10 cells significantly impaired the *in vivo* tumor growth (**Figure 5C**). *Zdhhc20* KO B16-F10 showed a significant decrease in both tumor volume and tumor weight (**Figure 5D-E**). We further repeated similar mouse experiments using the shCtrl and sh*Zdhhc20* B16-F10 cells which do not have the Cas9 protein expressed as a potential confounding neoantigen^46^ (**Figure S8F-G**). Similarly, *Zdhhc20* KD significantly decreased the tumor growth rate, tumor weight, and size (**Figure S8H-J**).

To confirm our hypothesis that the increased MHC-I level leads to a higher CD8^+^ T cell infiltration in the tumor microenvironment, we performed the same mice experiment and collected the tumor tissue on Day 10. We isolated the tumor infiltrated lymphocytes and quantified the immune cell population. Flow cytometry analysis showed increased tumor-infiltrating CD8^+^ T cells and CD45^+^ cells in the *Zdhhc20* KO B16-F10 tumor samples compared to WT tumor samples (**Figure 5F and Figure S9B**). We further confirmed the increased CD8^+^ T cells in the *Zdhhc20* KO B16-F10 tumor samples using immunohistology staining (CD8^+^ T cells were stained in Red, **Figure S9A**). Consistent with our findings in co-culture assays, we observed a significant increase of IFNγ^+^ or GZMB^+^ CD8^+^ T cells among the total tumor-infiltrating lymphocytes (TILs) when knocking out *Zdhhc20* (**Figure 5F**). Overall, the increase in tumor-infiltrating CD8^+^ T cells was in line with the reduced tumor burden when *Zdhhc20* was depleted.

In addition to melanoma, we also confirmed the effect of *Zdhhc20* in a mouse ovarian cancer model using the SO1 cell line (**Figure 5G-H**). *Zdhhc20* KD significantly increased the mouse survival rate compared to WT (**Figure 5I**).

We next wanted to further confirm that the effect of *ZdhhcC20* KO in inhibiting melanoma growth *in vivo* is dependent on CD8^+^ T cells. We first utilized the NSG mouse model which lacks mature T cells, B cells, and NK cells (**Figure 5J**)^47^. As expected, no significant difference was observed when comparing the growth curve, tumor weight, or tumor size between control and *Zdhhc20* KO B16-F10 cells, which suggests the *in vivo* growth difference is dependent on the tumor-immune interaction (**Figure 5K-M**). To determine whether the tumor-suppressive effect of *Zdhhc20* deficiency depends specifically on CD8^+^ T cells, we depleted CD8^+^ T cells in C57BL/6J mice using an anti-CD8α antibody prior to and following subcutaneous implantation of control or *Zdhhc20* KO B16-F10 cells (**Figure 5N**). Efficient CD8^+^ T-cell depletion was confirmed by flow cytometric analysis of both draining lymph nodes and peripheral blood mononuclear cells, where CD8^+^ T cells were nearly eliminated following anti-CD8α treatment (**Figure 5O and S9C**). Consistent with our previous findings, *Zdhhc20* KO significantly suppressed tumor growth in mice receiving the control antibody. In contrast, CD8^+^ T cell depletion largely abolished this growth inhibition, resulting in tumor growth kinetics and final tumor weights comparable to those of control tumors (**Figure 5P-R**). These data demonstrate that the antitumor effect elicited by *Zdhhc20* deficiency is predominantly mediated by CD8^+^ T cells.

### Loss of *ZDHHC20* enhances anti–PD-1 therapy

The expression level of MHC-I on tumor cells plays a critical role in determining the efficacy of therapies targeting the PD-1/PD-L1 axis. High MHC-I expression facilitates efficient presentation of tumor antigens to CD8 T cells, which are the primary effectors activated by PD-1/PD-L1 checkpoint blockade^48^. We therefore hypothesized that the upregulation of MHC-I presentation following *ZDHHC20* KO would enhance the therapeutic efficacy of PD-1/PD-L1 blockade. To test this, we analyzed two previously published melanoma patient datasets in which individuals received standard anti-PD-1 therapy using TIDE database^49–51^. Patients were stratified into two groups based on their *ZDHHC20* expression levels. Consistent with our hypothesis, patients with lower *ZDHHC20* expression exhibited significantly longer overall survival following anti-PD-1 treatment (**Figure 6A**). Since dichotomizing patients into high- and low-expression groups can create arbitrary distinctions between individuals immediately above and below the selected cutoff and may obscure the underlying biological relationship, we additionally performed a continuous analysis of the association between *ZDHHC20* expression and overall survival using two independent datasets available through cBioPortal and validated the similar findings (**Figure S10A-D**)^52–54^.

**Figure 6.**
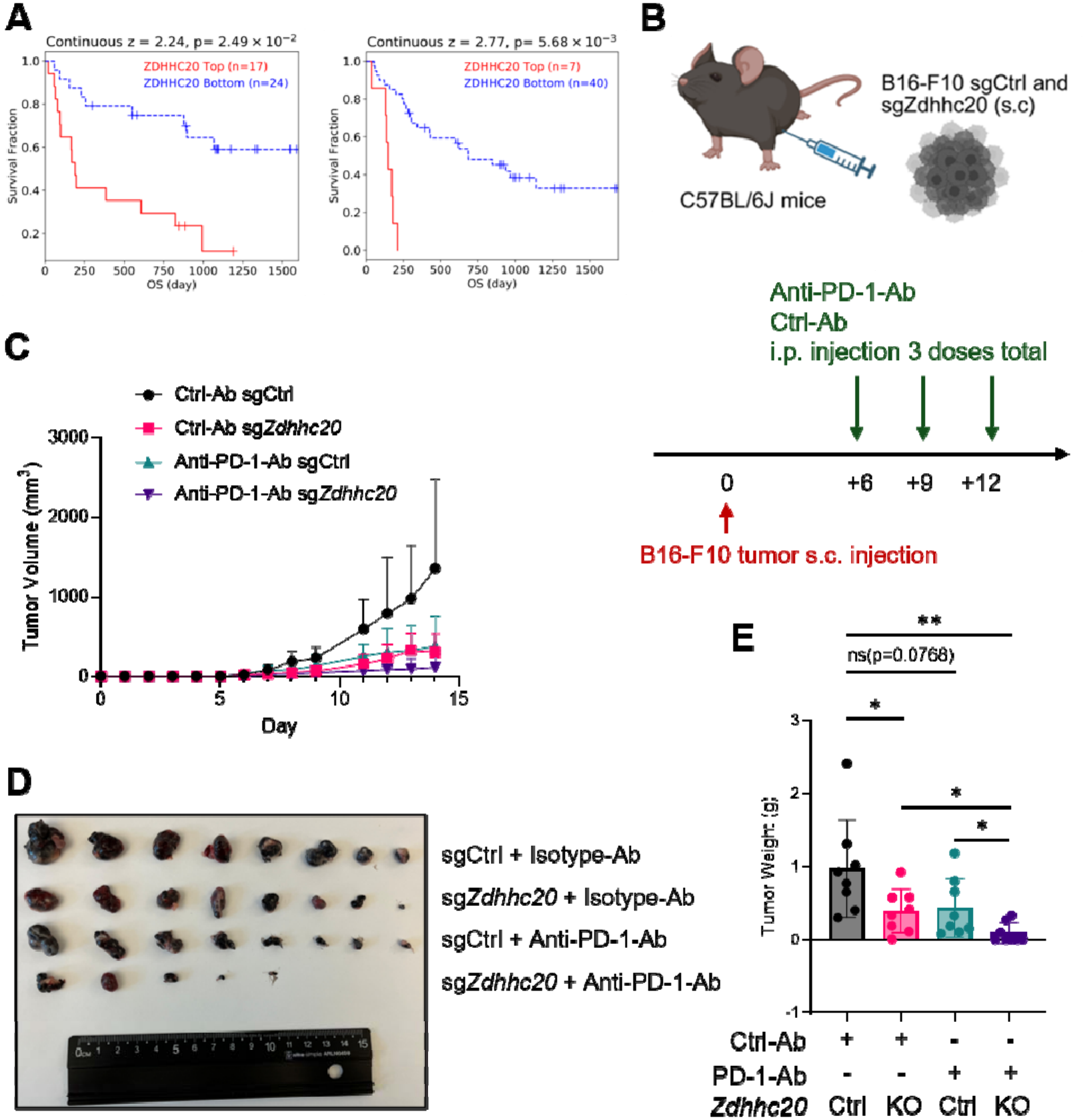
Loss of *Zdhhc20* enhances anti–PD-1 therapy. **A**. Survival analysis of patients receiving standard anti–PD-1 therapy with high and low *ZDHHC20* expression levels from two previous studies. ZDHHC20 mRNA expression (RNA-Seq TPM, log2 z-score-normalized relative to the cohort distribution) was used to stratify patients into high- and low-expression groups. The analysis is don using the TIDE server. **B-E**. B16-F10 subcutaneous tumor model using sgCtrl and sg*Zdhhc20* B16-F10 cells in C57BL/6J mice with anti-PD-1 co-treatment (N = 8 for each group). Anti–PD-1 and isotype control antibodies were injected through i.p. for three doses at Day 6, 9, and 12. Tumor growth curve (**C**), tumor image (**D**), and tumor weight (**E**) are shown. Statistical analysis: unpaired t test: (C, E).

We next sought to validate this finding using the B16-F10 melanoma mouse model in combination with anti-PD-1 therapy (**Figure 6B**). B16-F10 melanoma model typically has limited or modest responses to anti-PD-1 monotherapy^55^. In our hands, anti-PD-1 treatment alone produced a visible reduction in tumor growth in control B16-F10 tumors, although this effect did not reach statistical significance (*p* = 0.0768; **Figure 6C–E**). One possible explanation is the stable expression of bacterial Cas9 in our tumor cells, which has been reported to increase tumor immunogenicity in immunocompetent mice^46^. Importantly, combining *Zdhhc20* KO with anti-PD-1 therapy resulted in significantly greater tumor suppression, with five of eight tumors exhibiting little or no detectable growth (**Figure 6C–E**). These results support that targeting ZDHHC20 represents a promising strategy to improve the efficacy of immune checkpoint blockade therapy.

## Discussion

MHC-I plays a central role in cancer cell immune surveillance and tumor-intrinsic regulation of the antigen-presentation machinery constitutes a pivotal axis of immune evasion. It is well-documented that many cancers down-regulate MHC-I expression, through genetic deletion or mutation of heavy chains or β2-microglobulin, epigenetic silencing of transcriptional regulators such as NLRC5 or IRF1, post-translational degradation of MHC-I complexes, or defects in the APP pathway, thereby escaping CD8 T-cell recognition^22^. Clinically this translates into worse prognosis, reduced tumor-infiltrating lymphocytes, and resistance to ICB therapies^56^. Importantly, efforts to pharmacologically restore or up-regulate MHC-I have shown promise as an adjuvant to immunotherapy, thereby turning a ‘cold’ tumor into an immune-recognizable ‘hot’ one^57,58^.

Here, we identify a previously unrecognized ZDHHC20-TAP-MHC-I regulatory axis that plays critical roles in cancer immune evasion (**Figure 7**). ZDHHC20-mediated palmitoylation of the TAP1/2 complex promotes its ubiquitination by the ERAD E3 ligase RNF5/RNF185, leading to proteasomal degradation. The resulting reduction in the TAP complex suppresses MHC-I level. *ZDHHC20* KD in cancer cells increases MHC-I presentation and markedly enhances CD8 T-cell activation and cancer cell killing through the MHC-I-TCR axis *in vitro*. *In vivo*, loss of *ZDHHC20* significantly delays tumor growth in both melanoma and ovarian cancer models, accompanied by increased infiltration of activated CD8 T cells. Notably, this tumor-growth difference is abolished in immune-deficient NSG mice and in CD8 cytotoxic T cell-depleted mice, further underscoring the requirement for host immunity. Finally, genetic deletion of *ZDHHC20* enhances the therapeutic efficacy of anti-PD-1 immunotherapy.

**Figure 7.**
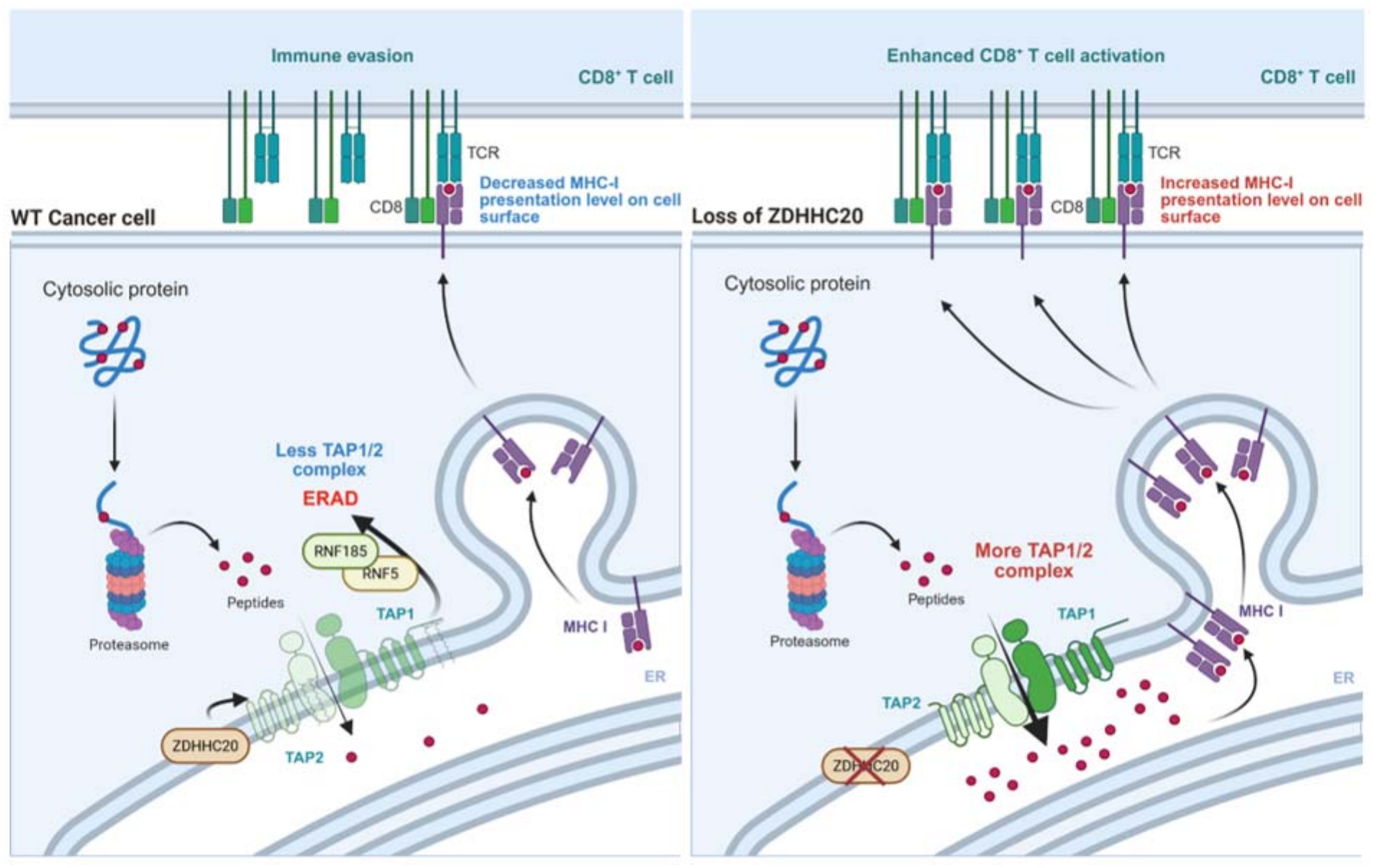
Graphic summary of ZDHHC20-TAP-MHC-I regulatory axis in cancer cells. In WT cancer cells, ZDHHC20-mediated palmitoylation of TAP1 and TAP2 promotes their RNF5/RNF185-dependent ERAD, reducing the abundance of the TAP1/2 complex. Consequently, fewer proteasome-generated peptides are transported into the ER for loading onto MHC-I molecules, resulting in decreased cell-surface MHC-I antigen presentation and facilitating immune evasion. Loss of ZDHHC20 stabilizes TAP1 and TAP2, increases TAP complex abundance and peptide transport into the ER, and enhance peptide–MHC-I presentation on the cancer cell surface. The increased recognition of peptide–MHC-I complexes by TCR promotes enhanced CD8 T-cell activation.

While protein cysteine palmitoylation has been known to regulate immune signaling and cancer progression,^9,10^ our study is the first to connect ZDHHC20 and protein cysteine palmitoylation to antigen presentation, a critical process in adaptive immunity. Regarding the role of ZDHHC20 in cancer progression, Zhang et al. reported that ZDHHC20-mediated palmitoylation of YTHDF3 prevents its autophagic degradation and stabilizes MYC mRNA, thereby promoting pancreatic cancer progression; correspondingly, *ZDHHC20* knockdown reduced the growth of human pancreatic cancer cell lines PANC-1 and AsPC-1 both *in vitro* and in immune-compromised mouse xenograft model^59^. Kharbanda et al. showed that loss of ZDHHC20-mediated palmitoylation of EGFR decreases cell proliferation *in vitro* and inhibits KRAS-mutant non-small cell lung cancer growth in mouse models^60^. More recently, Huang et al. found that ZDHHC20 palmitoylates and stabilizes GPX4, and that *ZDHHC20* depletion sensitizes the melanoma cell line A375 to ferroptosis^61^. Additionally, Tomić et al. reported that *ZDHHC20* deletion suppresses PDAC lung metastasis *in vivo* through effects on the innate immunity, mainly through NK cells^17^. Consistent with this finding, our analysis revealed a negative correlation between ZDHHC20 expression and NK cell infiltration in the tumor microenvironment, a broader role for ZDHHC20 in suppressing antitumor immunity (**Figure S11**). Our results in melanoma and ovarian cancer add a new dimension by revealing ZDHHC20’s previously unknown role in regulating the adaptive immune mechanism that may operate in parallel with these previously described tumor-intrinsic pathways or innate immune pathways.

ERAD is a well-established mechanism to ubiquitinate and degrade mis-folded or unassembled ER proteins. It is typically thought the E3 ligases involved in ERAD recognize exposed hydrophobic patches in mis-folded or unassembled ER proteins^36^. Our study reveals that palmitoylation may serve as a previously unknown degron recognized by ER–resident E3 ligases to degrade well folded proteins. We demonstrate that the ERAD-associated E3 ubiquitin ligases RNF5 and RNF185 are essential mediators of palmitoylation-dependent proteasomal degradation of TAP1 and TAP2. RNF5 and RNF185 share more than 70% sequence similarity and have previously been implicated in the ERAD of multiple substrates, including CFTR^41^, TAPBP^35^, CYP51A1^62^. A previous study connected ZDHHC6 Cys328 palmitoylation to ERAD. However, the specific E3 ubiquitin ligase responsible for recognizing palmitoylated ZDHHC6 was not identified^63^. To explore whether RNF5 and RNF185 could be the E3 that recognizes ZDHHC6 Cys328 palmitoylation, we performed AlphaFold3-based structural modeling. Interestingly, the model predicted a hydrophobic interaction between the Cys328-linked palmitoyl group and the phenylalanine-rich domains of both RNF5 and RNF185 (**Figure S12A-B**), like what we observed for TAP1 and TAP2. This further suggests that palmitoylation could serve as a degron recognized by the E3 ligases RNF5 and RNF185. Future efforts will likely identify more palmitoylation dependent substrates of those E3 ligases. However, the AlphaFold3 prediction may not be accurate and is only meant for hypothesis generation. The conformational transitions underlying ABC-transporter function and the distinct TAP states captured by viral inhibitors emphasize that potential recognition surfaces depend on transporter conformation and assembly context^25,27^. Accordingly, RNF5/RNF185 recruitment may involve contacts with a broader surface of the full-length TAP substrate, whose accessibility is influenced by heterodimerization, PLC association and the surrounding membrane, rather than the palmitoylated cysteine alone. Palmitoylation could contribute directly to this interface or indirectly alter local membrane interactions and substrate conformation. The AlphaFold3 models therefore may provide hypothesis consistent with the biochemical observations, but do not independently establish direct recognition of the palmitoyl chain. It also remains unresolved whether RNF5/RNF185 preferentially engage PLC-associated TAP or unassembled or partially assembled intermediates. Distinguishing these possibilities will require direct interaction mapping and structural or reconstitution studies that preserve the relevant membrane and assembly context.

The effect of palmitoylation on protein stability is highly substrate dependent. For example, ZDHHC3-mediated palmitoylation of PD-L1 at Cys272 inhibits its ubiquitination and lysosomal degradation, thereby stabilizing PD-L1 at the cell surface^14^. Likewise, palmitoylation has been reported to regulate the stability and function of several other proteins, including NLRP3^64,65^, SCRIB^66^, Fas^67^ through distinct molecular mechanisms. In contrast, our study identifies TAP1 and TAP2 as substrates whose palmitoylation promotes ER-associated degradation, highlighting that palmitoylation can either stabilize or destabilize proteins depending on the molecular context and the downstream protein quality-control machinery engaged.

Understanding how the ZDHHC20-TAP-MHC-I axis is dysregulated during tumorigenesis will be important for defining the patient populations most likely to benefit from therapeutic targeting of ZDHHC20. Interestingly, Zhang et al. demonstrated that ZDHHC20 expression is induced by oncogenic KRAS signaling, linking activation of a major oncogenic pathway to increased ZDHHC20 abundance^59^. These findings raise the possibility that KRAS-driven cancers may exploit ZDHHC20-mediated TAP degradation to impair antigen presentation and facilitate immune evasion. The expression profile of ZDHHC20 suggests that it is broadly upregulated across 23 tumor types, without an obvious lineage-specific expression pattern (**Figure S1E and Figure S2**)^23^. However, these analyses are primarily based on transcriptomic datasets. Future studies incorporating more protein-level based measurements in clinical specimens will be essential to better define the regulation of ZDHHC20 in human cancers and to determine whether its protein abundance correlates with genetic alterations, oncogenic signaling, or immune evasion. The clinical relevance of this ZDHHC20-TAP-MHC-I regulatory axis is further underscored by analysis of human patient data. Nevertheless, further validation in independent clinical cohorts will be important to define the clinical significance of the ZDHHC20-TAP-MHC-I axis in antitumor immunity and response to immune ICB.

ICB therapy has revolutionized cancer therapy by restoring antitumor T cell responses, particularly through antibodies targeting PD-1, PD-L1, and CTLA-4^68–70^. Despite durable clinical responses in a subset of patients, the majority of tumors exhibit primary or acquired resistance to ICB^71^. A central challenge is the prevalence of “immune cold” tumors, which are characterized by low CD8 T cell infiltration, impaired antigen presentation, and reduced MHC-I surface expression, thereby limiting effective T cell priming and cytotoxic activity. Converting ‘immune cold’ tumors into ‘immune hot’ tumors with robust antigen presentation and T cell recruitment has therefore emerged as a major therapeutic goal. Our findings identified ZDHHC20 as a critical negative regulator of tumor antigen presentation through palmitoylation-dependent degradation of the TAP1/2 complex. Targeting ZDHHC20 could thus restore antigen presentation, enhance intratumoral CD8 T cell accumulation, and sensitize immune–excluded tumors to ICB therapy. These results highlight ZDHHC20 inhibition as a potential strategy to improve the efficacy of ICB therapies.

## Materials and Methods

### Cell culture, transfection and lentivirus transduction

Human HEK293T, A549, A375 cells, Mouse B16-F10, IE9mp1, SO1, RAW264.7 were cultured in DMEM (Gibco) media supplemented with 10% (v/v) heat-inactivated fetal bovine serum (FBS, Gibco). Mouse DC2.4 cells were cultured with RPMI-1640 (Gibco) media supplemented with 10% (v/v) FBS, 1x GlutaMAX (Gibco), 1x non-essential amino acids (TMS-001-C, Sigma), 1x HEPES Buffer Solution (TMS003-C, Sigma) and 0.0054x β-Mercaptoethanol. For transient overexpression in HEK293T cells, the transfection was performed using polyethylenimine (PEI, 24765, Polysciences). For siRNA mediated transient KD, the transfection was performed using the Lipofectamine RNAiMAX (13778150, Thermo) according to the manufacturer’s instructions. For the rescue experiments in A549 cells, the transfection was performed using the NanoFect Transfection Reagent (NF100, ALSTEM) according to the manufacturer’s instructions.

sgRNA, shRNA or stable overexpression (pCDH and pLenti vectors) plasmids (4 μg) were transfected into HEK293T cells together with the psPAX2 (2.8 μg) and pMD2.G (1.2 μg) packaging plasmids using PEI. The lentivirus-containing supernatant was collected at 48 h following transfection. Cells were transduced with lentivirus supplemented with 8 μg/mL Polybrene for 24 h and recovered in complete medium for 24 h. The cells were then subjected to 1∼5 μg/mL puromycin (H-270-1, Goldbio) or 20 μg/mL blasticidin (ant-bl, InvivoGen) or 100 μg/mL hygromycin (ant-hg-1, InvivoGen) selection for 48∼72 h. lentiCas9-Blast was a gift from Feng Zhang (Addgene plasmid # 52962 ; http://n2t.net/addgene:52962 ; RRID:Addgene_52962). pT2/SV-Venus-SIINFEKL was a gift from Jon Yewdell (Addgene plasmid # 111624 ; http://n2t.net/addgene:111624 ; RRID:Addgene_111624). All the other key plasmids, primers and reagents used are listed in the **supplementary file**.

### Cell viability measurement

Cells were seeded in 96-well plates at densities of 10,000, 20,000, or 50,000 cells per well. After 72 hours of incubation at 37 °C, CellTiterGlo 2.0 assay (Promega) was performed according to the manufacturer’s instructions. Cell viability (%) was calculated by normalizing to the WT control cells.

### Cloning and mutagenesis

pEGFP-ZDHHC20 and empty control plasmid were a kind gift from Dr. Maurine Linder. The TAP1 and TAP2 sequences are cloned from the Human TAP1 cDNA ORF (Sino biological, HG17575-UT) and Human TAP2 cDNA ORF (Sino biological, HG17653-U), respectively. pCMV3-TAP1-C-Flag and pCMV3-TAP2-C-Flag were generated using HiFi DNA Assembly kit (NEB, E5520). Point mutations of plasmids were generated by Q5 site directed mutagenesis (NEB, E0554S). pLenti-TAP2 was ordered from VectorBuilder. Primers are listed in the supplementary file.

### Immunoprecipitation

Cells transfected with indicated plasmids for 24 h were collected and lysed in RIPA buffer supplemented with protease inhibitor cocktail and nuclease. After centrifugation at 16,000 × g for 10 min at 4 °C, the supernatant was collected. 20 μL of anti-Flag agarose beads were washed using Flag-IP washing buffer three times and then added into each sample. Following rocking at 4 °C for 3 h, the lysates were centrifugated at 2,000 × g for 2 min at 4 °C to spin down the beads. The beads were then washed using Flag-IP washing buffer three times and directly eluted by adding 12 μL of 6× SDS loading buffer with boiling at 95 °C for 10 min. The eluted samples were resolved by SDS-PAGE and analyzed by western blotting.

For TAP1/2 and TAPBP ubiquitination level and RNF5/185 interaction level analysis, the samples were treated with 20 μM of Mg132 (MCE, HY-13259) for 6 h before collection.

### Click chemistry and in-gel fluorescence detection (Alk14 labeling)

Alk14 labeling assays were performed as described previously^72^. Cells were treated with 100 μM palmitic acid analogue Alkyne 14 (Alk14) for 6 h, then collected and lysed in RIPA buffer supplemented with protease inhibitor cocktail and nuclease. After centrifugation at 16,000 × g for 10 min at 4 °C, the supernatant was collected. The target protein was purified using anti-Flag agarose beads following the immunoprecipitation procedure, which were subsequently suspended in 20 μL of Flag-IP washing buffer. Click chemistry reagents were added to the beads in the following order: 1 μL of 4 mM TAMRA azide (47130, Lumiprobe), 1 μL of 10 mM tris[(1-benzyl-1H-1,2,3-triazol-4-yl)methyl]amine (TBTA; T2993, Tcichemicals), 1 μL of 40 mM CuSO, and 1 μL of 40 mM tris(2-carboxyethyl)phosphine hydrochloride (TCEP HCl; 580560, Millipore). The mixtures were mixed thoroughly and incubated for 30 min at RT in the dark. Subsequently, 12 μL of 6 × SDS loading buffer was added, and the samples were heated at 95 °C for 10 min. Proteins were resolved by SDS–PAGE, and rhodamine fluorescence was detected using Chemidoc.

### Acyl-biotin exchange (ABE)

ABE assays were performed as described previously^73^. Samples were suspended in 1 ml of lysis buffer (100 mM Tris-HCl, pH 7.2; 5 mM EDTA; 150 mM NaCl; 2.5% SDS; protease inhibitor cocktail) supplemented with 50 mM N-ethylmaleimide (NEM; E3876, Sigma) and 50 U ml ¹ nuclease (88700, Thermo Fisher). The samples were solubilized at RT for 1 h with gentle mixing. Protein concentrations in the supernatants were determined using BCA assay. Samples were adjusted to 1 mg/mL in lysis buffer to a final volume of 1.1 mL. Subsequently, 12 μL of 2,3-dimethyl-1,3-butadiene (final concentration ∼100 mM) were added to each sample and incubated at RT for 1 h with mixing. One tenth volume of chloroform was then added to remove residual 2,3-dimethyl-1,3-butadiene. Samples were then divided equally into two portions and incubated with either 0.5 ml of 1 M hydroxylamine or 1 M NaCl. Then, 100 μL of 1 mM biotin-HPDP solution were added to both aliquots and incubated for 1 h at RT. Following incubation, proteins were precipitated with chloroform/methanol/water (v/v 1:4:3) and resuspended in 200 μL of resuspension buffer (100 mM Tris-HCl, pH 7.2; 2% SDS; 8 M urea; 5 mM EDTA). A 20 μL aliquot was saved as the loading control, while the remaining 180 μL was diluted 1:10 with PBS and incubated with 20 μL of streptavidin beads for 1 h at RT with shaking. Beads were washed three times with PBS containing 1% SDS. Both bead-bound samples and loading controls were mixed with SDS loading buffer and heated at 95 °C for 10 min. Proteins were resolved by SDS–PAGE and analyzed by western blotting.

### SIINFEKL-based MHC-I presentation

The SIINFEKL-based MHC-I presentation assay was performed following a previously published method^21^. B16-F10 cells were transfected using Lipofectamine 3000 Transfection Reagent to overexpress either GFP-EV or Venus-SIINFEKEL for 24 h. Cells were then disassociated using the Cellstripper (Non-enzymatic cell dissociation solution, 25-056-CI, Corning) and labeled using the APC anti-mouse H-2Kb bound to SIINFEKL Antibody (141605, Biolegend). The green fluorescence positive cell population was first gated to sort out the successfully transfected cells. Then the H-2Kb-SIINFEKEL^+^ cell population was quantified using the Attune NxT Flow Cytometer.

### Mice experiment

C57BL/6 mice and NSG mice were purchased from the Jackson Laboratories. All animal experiments were carried out in accordance with the guidelines of the National Advisory Committee on Laboratory Animal Research and the University of Chicago Institutional Animal Care and Use Committee.

Subcutaneous xenograft tumor models were established by inoculating B16-F10 cells into the flank of C57BL/6J mice (8-10 weeks old, male). Tumor growth was monitored daily from day 6, with volume calculated as 0.5□×□length□×□width². Mice were sacrificed on day 12-15 (B16-F10) once the average tumor volume reached 1000 mm^3^ and the tumors were extracted for tumor weight measurement and photo. For the tumor-infiltrating lymphocytes analysis, mice were sacrificed on day 10 and the tumors were extracted for flow cytometry analysis. To evaluate the impact of *Zdhhc20* deficiency on tumor growth in vivo, sgCtrl (or shCtrl) and sg*Zdhhc20* (or sh*Zdhhc20*) B16-F10 (1□×□10^5^ in 100 µL of sterile PBS) cells were subcutaneously inoculated into C57BL/6J mice. To investigate whether the effect of *Zdhhc20* is through tumor-host immune interaction, the same procedures were repeated using the NSG mice (8-10 weeks old, male, 005557, The Jackson Laboratory). For the CD8^+^ T cell depletion model, mice were randomized into two groups: one treated with 200 μg isotype control (clone LTF-2, BE0090, BioXCell) and the other with 200 μg anti-CD8α antibody (clone 2.43, BE0061, BioXCell) via i.p. injection on days −3, 0, 3 and 8 (four doses total). On day 0, B16-F10 cells (1□×□10^5^) were subcutaneously injected. To assess the combined efficacy of *Zdhhc20* deficiency and anti-PD-1 antibody treatment. Mice were randomized into four groups: (A) sgCtrl B16-F10 with isotype control (clone 2A3, BE-0089, BioXCell); (B) sg*Zdhhc20* B16-F10 with isotype control; (C) sgCtrl B16-F10 with anti-PD-1 antibody (clone RMP1-14, BE-0146, BioXCell); and (D) sg*Zdhhc20* B16-F10 with isotype control with anti-PD-1 antibody. sgCtrl and sgZdhhc20 B16-F10 cells (1□×□10^5^) were subcutaneously injected into group A/B and Group C/D, respectively. Isotype control and anti-PD-1 antibody were i.p. injected (200 μg per injection) on days 6, 9, and 12.

Female C57BL/6 mice (8 weeks old) were used for the SO1 intraperitoneal ovarian tumor model^74^. A total of 1 × 10^6^ SO1 cells suspended in 100 µL of sterile PBS were injected intraperitoneally into each mouse. Following tumor implantation, mice were monitored daily for signs of distress, body weight loss, and overall health status. Survival was recorded from the day of tumor inoculation until the experimental endpoint, defined by humane criteria in accordance with institutional animal care guidelines.

### Flow cytometry assay

For cancer cells, samples were detached using Cellstripper buffer (Corning, 25-056-CI) and stained with the indicated antibodies according to the manufacturer’s instructions. For lymphocytes, cells were first treated with TruStain FcX™ (anti-mouse CD16/32) antibody to block Fc receptors, followed by staining with surface protein markers. Cells were then labeled with the LIVE/DEAD™ Fixable Aqua Dead Cell Stain Kit to assess viability. For intracellular protein detection, cells were processed using the Intracellular Fixation & Permeabilization Buffer Set (Thermo Fisher, 88-8824-00) and subsequently stained with fluorescently labeled intracellular antibodies. To assess CD8 T cell effector function, TILs were stimulated for 6h at 37 °C with Cell Activation Cocktail containing Brefeldin A (BioLegend, 423304) prior to antibody staining. Flow cytometry data were acquired on either Attune NxT (Thermo Fisher), Cytek Aurora (Cytek Biosciences) or Quanteon (Agilent) flow cytometer and analyzed using FlowJo v10.0. All the antibodies used are shown in supplementary files.

### Reverse transcription and quantitative PCR (RT-qPCR)

Total mRNA was extracted using the E.Z.N.A.® Total RNA Kit I from cells (R6834, Omega Bio-tek). cDNA was then obtained by High-Capacity cDNA Reverse Transcription Kit (4368813, Thermo Fisher). qPCR was performed using SYBR Green PCR Master Mix (4309155, Thermo Fisher) with QuantStudio™ 7 Flex Real-Time PCR System. Relative expression of genes was calculated using the ΔΔCq method and normalized to the control gene GAPDH. Primer sequences are listed in the supplementary files.

### Histology

For histology sample preparations, mouse tumor tissues were fixed by 10% (v/v) neutral buffered formalin solution, embedded in paraffin, sectioned and stained with CD8a antibody (CST, 98941). Slides were imaged using the CRi Pannoramic MIDI 20x Whole Slide Scanner. The images were analyzed using QuPath v0.6.0.

### Isolation of tumor-infiltrating lymphocytes

Mouse tumors were extracted and kept in RPMI media to prevent dehydration and TIL loss. Tumor samples were then minced into small pieces using a scalpel and transferred to a 15 mL tube containing 5 mL tumor digestion media (500 μL Collagenase/Hyaluronidase solution, 750 μL 1 mg/mL DNase I solution and 3.75 mL RPMI 1640 medium). The digestion mixtures were then incubated with shaking for 40 min at 37 °C. The digested tumor tissues were transferred into a 70 μm mesh nylon strainer on a 50-mL conical tube and meshed using the rubber end of a syringe plunger. The collected tumor tissues were centrifugated at 300 g for 10 min at RT to collect the single cell suspensions. 10 mL of ammonium chloride solution was then added to lyse the red blood cells at RT for 5 min. After washing with 10 mL of PBS, the cell solutions were centrifugated at 300 g for 10 min at 4 °C. TILs were isolated using Ficoll-Paque Plus (GE Healthcare) density gradient centrifugation for 40 min at 300 g with the brake set on low. The layer of mononuclear cells was transferred to a sterile 15-mL conical tube and further washed with RPMI medium to obtain the TILs.

### In vitro assays for T-cell recognition

HLA-A*0201-resticted NY-ESO-1 antigen-specific TCR gene-transduced and untransduced T cells were generated as previously described^42^. T cells (1 × 10^5^) were stimulated with A375 (HLA-A*0201^+^NY-ESO-1^+^) transfected with shCtrl or sh*ZDHHC20* (0.5 × 10^5^) for 6 hours in the presence of 5 µg/ml monensin (Sigma) in a 96-well round bottom plate. After the incubation, cells were harvested and stained for CD8 BV421 (clone SK1, BioLegend), fixed and permeabilized using BD Cytofix/Cytoperm kit (BD Biosciences), followed by intracellular staining for IFN-γ APC (clone B27, BioLegend). The samples were acquired by an Attune NxT flow cytometer (Invitrogen) and the FCS files were analyzed using FlowJo software (Version 10.8.0).

To analyze IFN-γ release from T cells against target cells, T cells (1 × 10^5^) and A375 (0.5 × 10^5^) were cocultured for 24 hours in a 96-well round bottom plate. The culture supernatant was harvested and IFN-γ levels were measured using a human IFN-γ ELISA kit (Invitrogen) according to the manufacturer’s instructions.

To determine T-cell mediated apoptotic cell death of A375, A375 (1 × 10^5^) was seeded to a 24-well plate and T cells (2 × 10^5^) were added when A375 cells were attached to the plate. After 24-hour incubation, all cells were harvested with 0.25% trypsin/EDTA solution, stained with propidium iodide (PI) solution and anti-annexin V FITC antibody (BioLegend) in annexin V binding buffer (BioLegend). The samples were analyzed by flow cytometry.

For in vitro murine T-cell activation analysis, splenocytes (1 × 10^5^) obtained from RAG-1^-/-^ OT-1 mice were cocultured with ovalbumin (OVA)-expressing IE9mp1^75^ transfected with shCtrl or sh*Zdhhc20* at different E:T ratios in a 96-well round bottom plate for 24 hours. The cells were harvested, stained with anti-CD8 AF488 (clone 53-6.7, BioLegend) and anti-CD69 PE/Cy5 (clone H1.2F3, BioLegend) antibodies, and analyzed by flow cytometry.

### AlphaFold3 prediction

AlphaFold3 structural predictions were performed using the AlphaFold Server^38^. For the curated panel of 15 E3 ligases involved in ERAD, each ligase was modeled for potential interaction with either unmodified or palmitoylated TAP1 and TAP2 individually. All predicted complexes were manually examined to assess (A) conformational changes in binding associated with palmitoylation and (B) the spatial distance between the palmitoyl group and the dimer interface. The 15 E3 ligases analyzed were: SYVN1 (HRD1), AMFR (GP78), MARCHF6 (TEB4), RNF5 (RMA1), TRC8, TMEM129, RNF185, UBE3C, TRIM13, RNF103, RNFT1, CGRRF1, RNF19B, PARK2 (PARKIN), and STUB1 (CHIP). Those previously known as interacted E3 ligases are also modeled as a complex with either TAP1 or TAP2 individually.

### Data mining and bioinformatics analysis

Overall survival (OS) analysis of melanoma patients receiving anti–PD-1 therapy (**Figure 6A**) was performed using the TIDE database^51^. OS analysis of extra melanoma patients receiving ICB (**Figure S10**) was performed using the cBioPortal^54^. Correlation analysis of CD8 T-cell infiltration level and *ZDHHC20* expression (**Figure 1A-B and Figure S11**) was performed using CIBERSORT^76^. In brief, the expression levels of ZDHHC family members were correlated with CD8 T-cell infiltration across 40 TCGA cancer types. For each *ZDHHC* gene, the number of cancer types showing a significant correlation (either negatively or positively) was quantified and ranked accordingly. Correlation analysis of *CCL5/CD3D* level and *ZDHHC20* expression (**Figure S8B-C**) was performed using Timer2.0 database^15^.

### Statistical analysis

For biological experiment, data were analyzed by Prism v.10 software (GraphPad) using two-tailed unpaired Student’s t-tests, one-way ANOVA or two-way ANOVA, as indicated in the figure legends. The Mantel–Cox test was used for comparing mouse survival curves.

## Acknowledgement

We thank The University of Chicago Human Tissue Resource Center (RRID:SCR_019199), especially Terri Li, for their assistance with histology sample preparation. We thank Dr. Jing Chen, Dr. Megan Borregard, Dr. Jiacheng Li, Dr. Zhicheng Xie, and Yuxin Zhuang for their helpful discussions. We thank Dr. Maurine Linder for providing the pEGFP-ZDHHC20-WT plasmids. We thank Dr. Shaoyi Jiang for providing DC2.4 cells. We thank Bryan Arpi for providing the HEK293T Cas9 cells. KO, JM, JL, FQ, K.G. are supported by NCI P30 grant NCI P30CA014599-50 to the University of Chicago Comprehensive Cancer Center. Flow cytometry was performed at the Cytometry and Antibody Technology Facility at the University of Chicago, which receives financial support from the Cancer Center Support Grant (P30CA014599). RRID: SCR_017760. We thank the University of Chicago Proteomics Platform (RRID:SCR_022928).

## Conflict of Interest Statement

K.O. is a co-founder of Tactiva Therapeutics. Other authors have no conflict of interest to declare.

---

**Figure S1.**
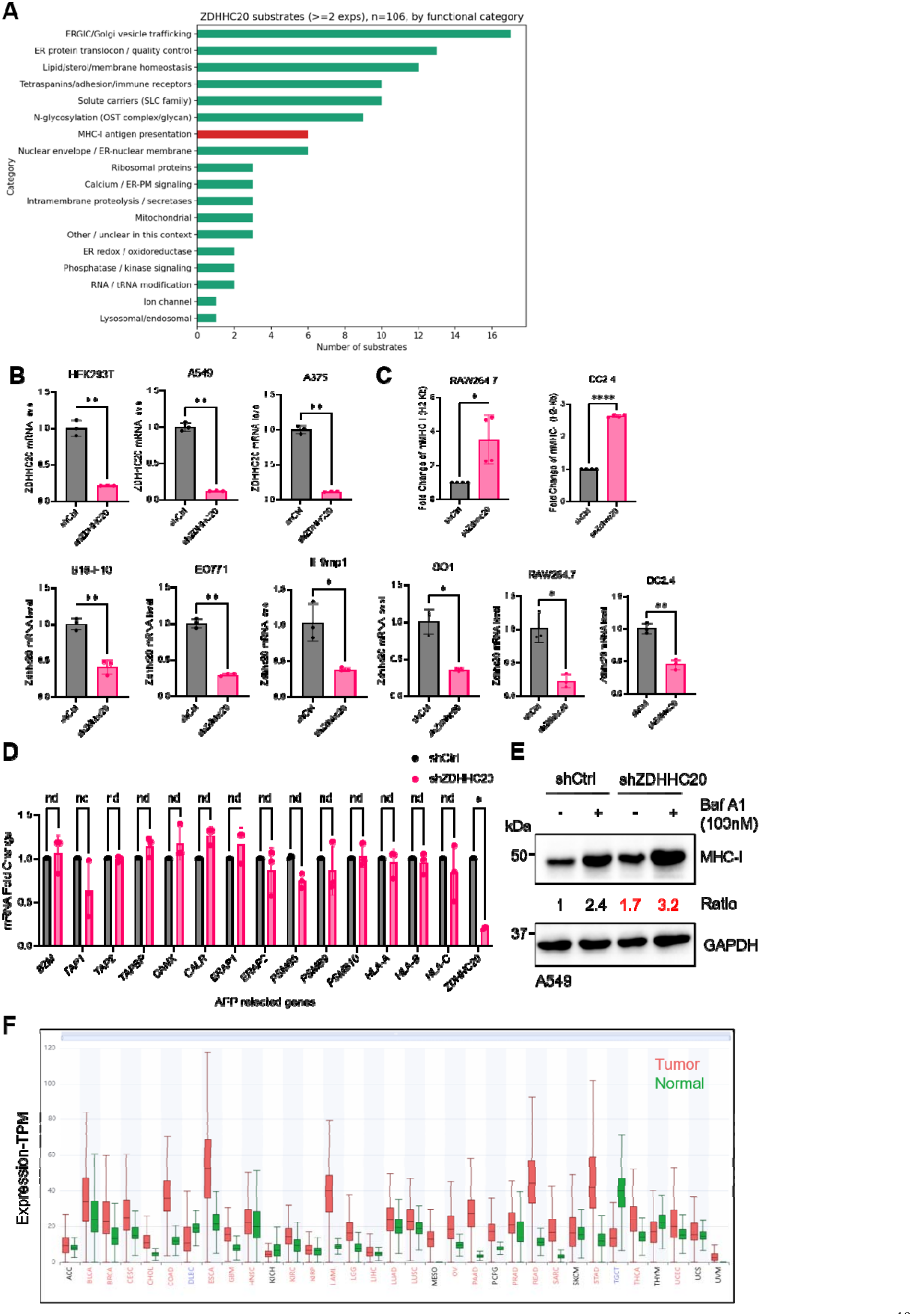
**A**. Analysis of ZDHHC20 substrates using Bump-hole strategy by functional categories^19^. **B**. qPCR validation of *ZDHHC20* knockdown in different cell lines. **C**. Zdhhc20 knockdown in RAW264.7 and DC2.4 cells increases MHC-I levels. **D**. mRNA fold-changes of antigen presentation-related genes in shCtrl and sh*ZDHHC20* HEK293T cells. **E**. MHC-I levels in Ctrl and ZDHHC20 knockdown A549 cells treated with or without Bafilomycin A1 (100 nM, overnight). **F**. *ZDHHC20* gene-expression levels in tumor versus normal tissues. Tissues with significantly higher expression are highlighted with red label, significantly lower expression in blue label, and those with no significant difference in black label. Data obtained from GEPIA3.0. Statistical analysis: unpaired t test: (B, D); paired t test: (C); multiple unpaired test: (D). Sample sizes: (B, C, D) n = 3. Image in E shows representative results from two independent experiments.

**Figure S2.**
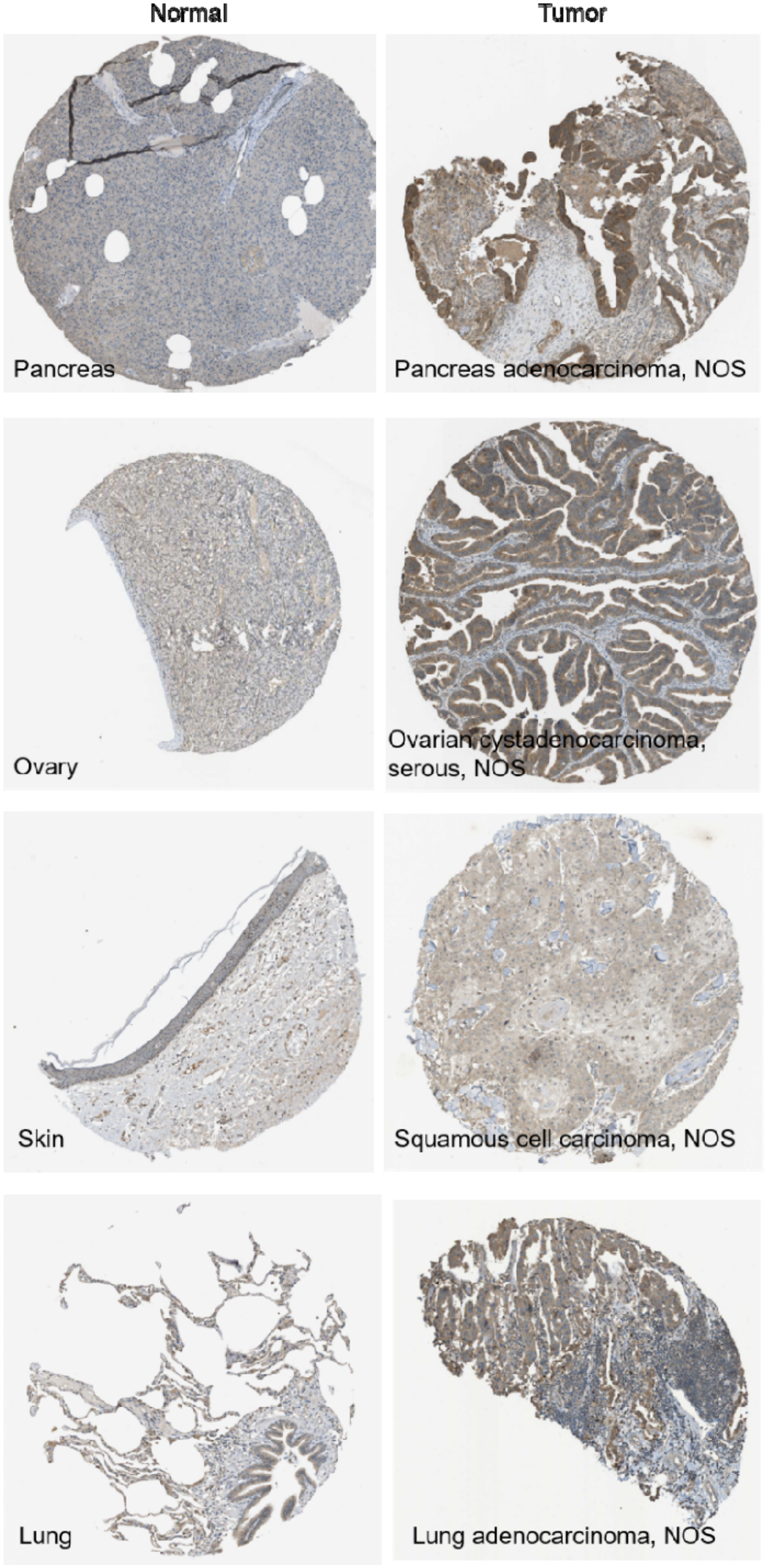
ZDHHC20 protein expression is elevated in multiple human cancers. Representative immunohistochemical (IHC) staining images of ZDHHC20 protein expression in normal tissues and their corresponding tumor tissues obtained from the Human Protein Atlas^24^. Representative tissue pairs includ pancreas and pancreatic adenocarcinoma, ovary and ovarian serous adenocarcinoma, skin and cutaneous squamous cell carcinoma, and lung and lung adenocarcinoma. Increased ZDHHC20 level is observed in tumor tissues compared with the corresponding normal tissues, supporting the broad upregulation of ZDHHC20 across multiple cancer types (Antibody HPA014702).

**Figure S3.**
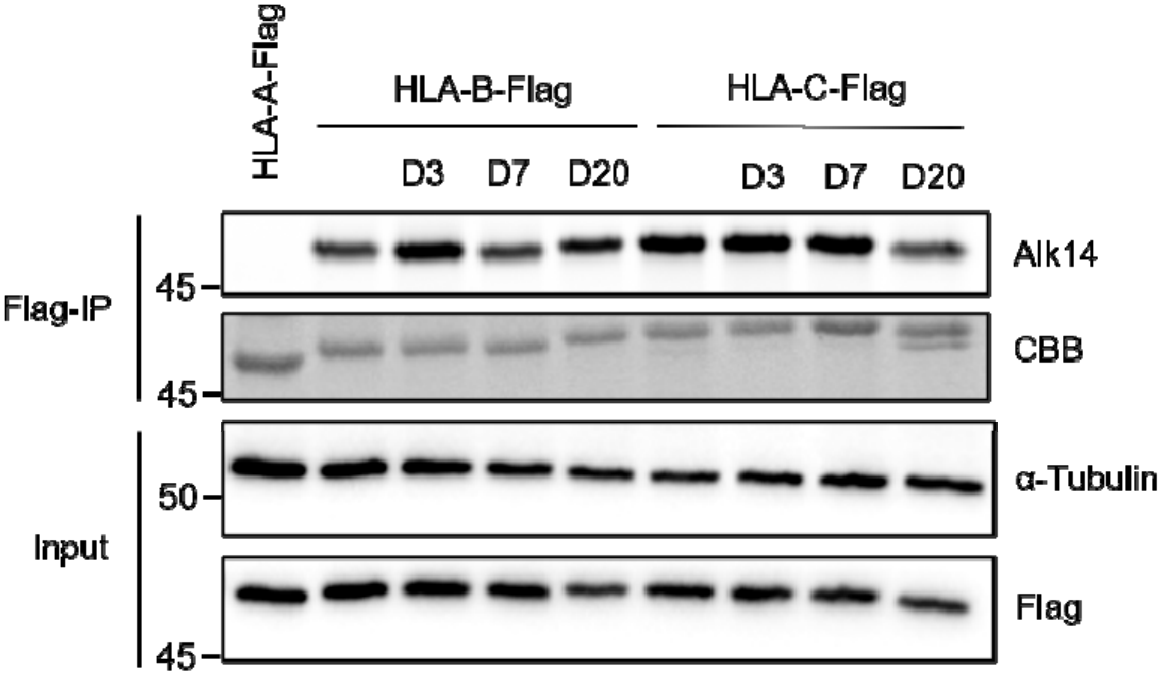
HLA-B and C are not the substrates of ZDHHC20. HEK293T cells expressing Flag-tagged HLA-A or HLA-B, HLA-C with ZDHHC3/7/20 were metabolically labeled with Alk14.

**Figure S4.**
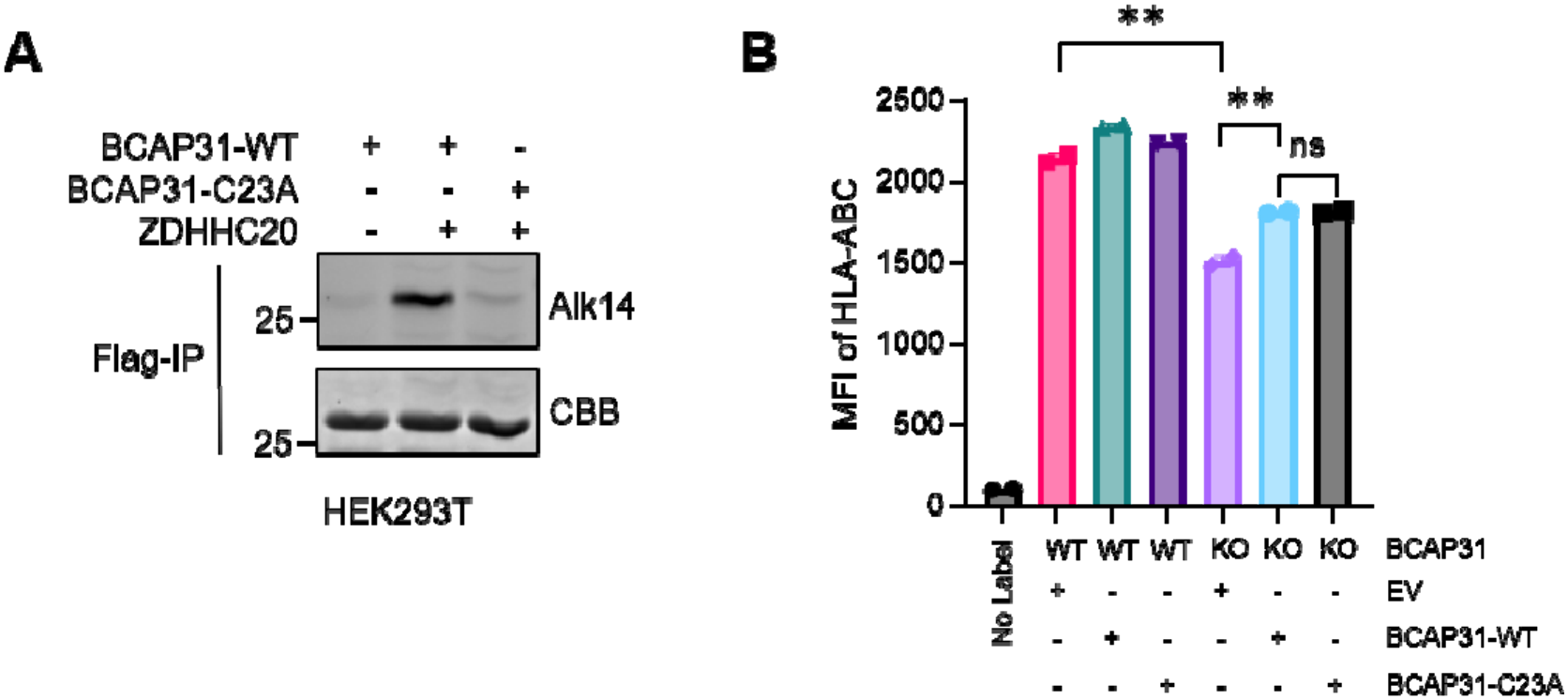
BCAP31 is a substrate of ZDHHC20, but its palmitoylation does not regulate MHC-I expression. **A**. HEK293T cells expressing Flag-tagged BCAP31-WT or the palmitoylation-deficient mutant BCAP31-C23A together with GFP-ZDHHC20 were metabolically labeled with Alk14. **B**. Flow cytometric analysis of surface HLA-ABC expression in WT and *BCAP31* KO HEK293T cells reconstituted with EV, BCAP31-WT, or BCAP31-C23A. *BCAP31* KO reduced surface HLA-ABC expression, whereas re-expression of BCAP31-WT or the palmitoylation-deficient C23A mutant restored HLA-ABC expression to a similar extent, indicating that BCAP31 palmitoylation is dispensable for MHC-I surface expression. Image in A shows representative results from two independent experiments. Sample sizes: (B) N = 2.

**Figure S5.**
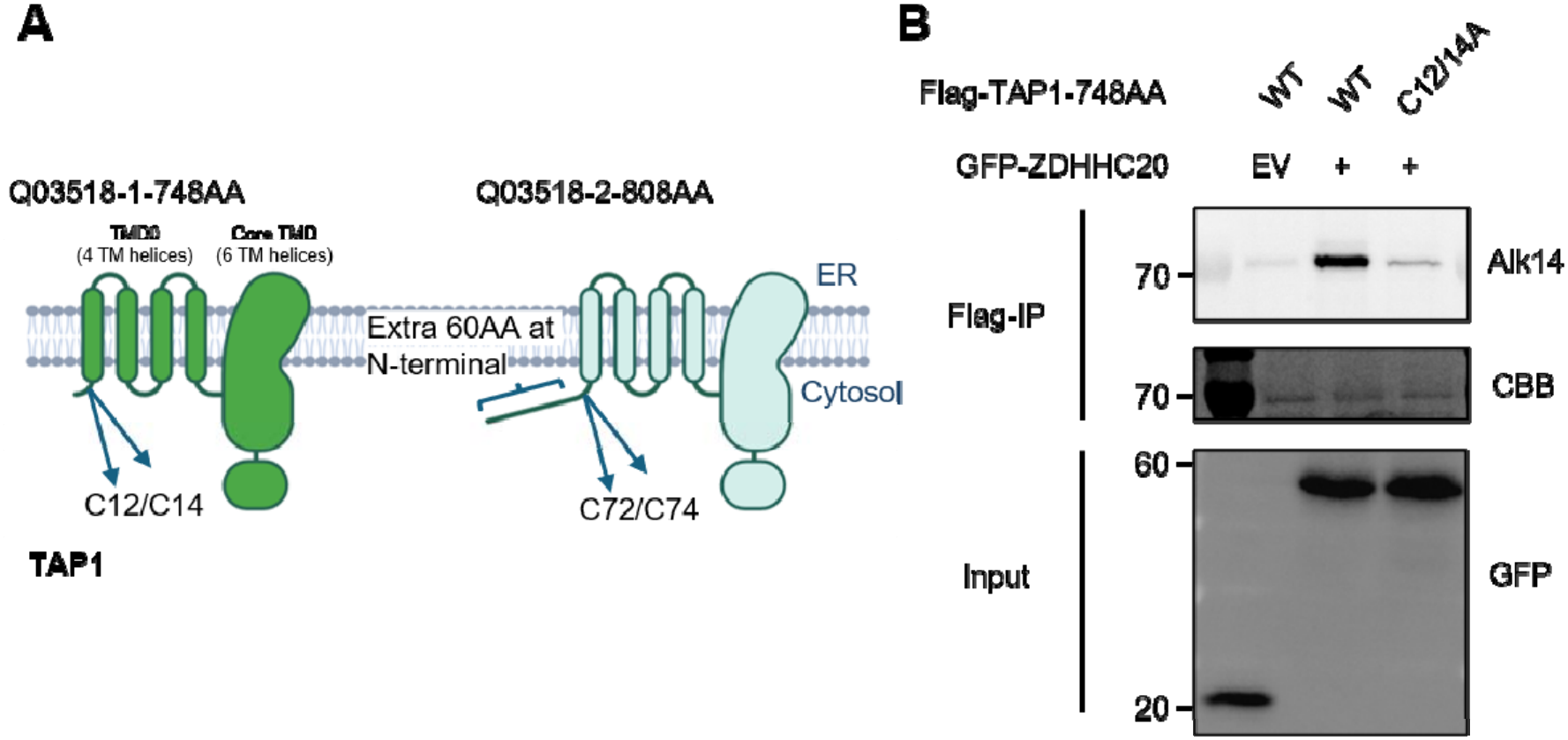
ZDHHC20 promotes TAP1 palmitoylation at Cys12 and Cys14 in the 748-amino-acid isoform. (**A**) Schematic representation of the two human TAP1 isoforms, Q03518-1 (748 amino acids) and Q03518-2 (808 amino acids). TAP1 contains an N-terminal transmembrane domain (TMD0; four transmembrane helices) and a core transmembrane domain (six transmembrane helices). The additional 60 N-terminal residues in Q03518-2 shift the numbering of Cys12 and Cys14 in Q03518-1 to Cys72 and Cys74, respectively. ER, endoplasmic reticulum. (**B**) Alk14 labeling of FLAG-tagged TAP1 (748-amino-acid isoform), either WT or the C12/14A double mutant, co-expressed with GFP–ZDHHC20 or GFP-EV. The C12/14A mutation markedly reduced ZDHHC20-dependent Alk14 labeling of TAP1.

**Figure S6.**
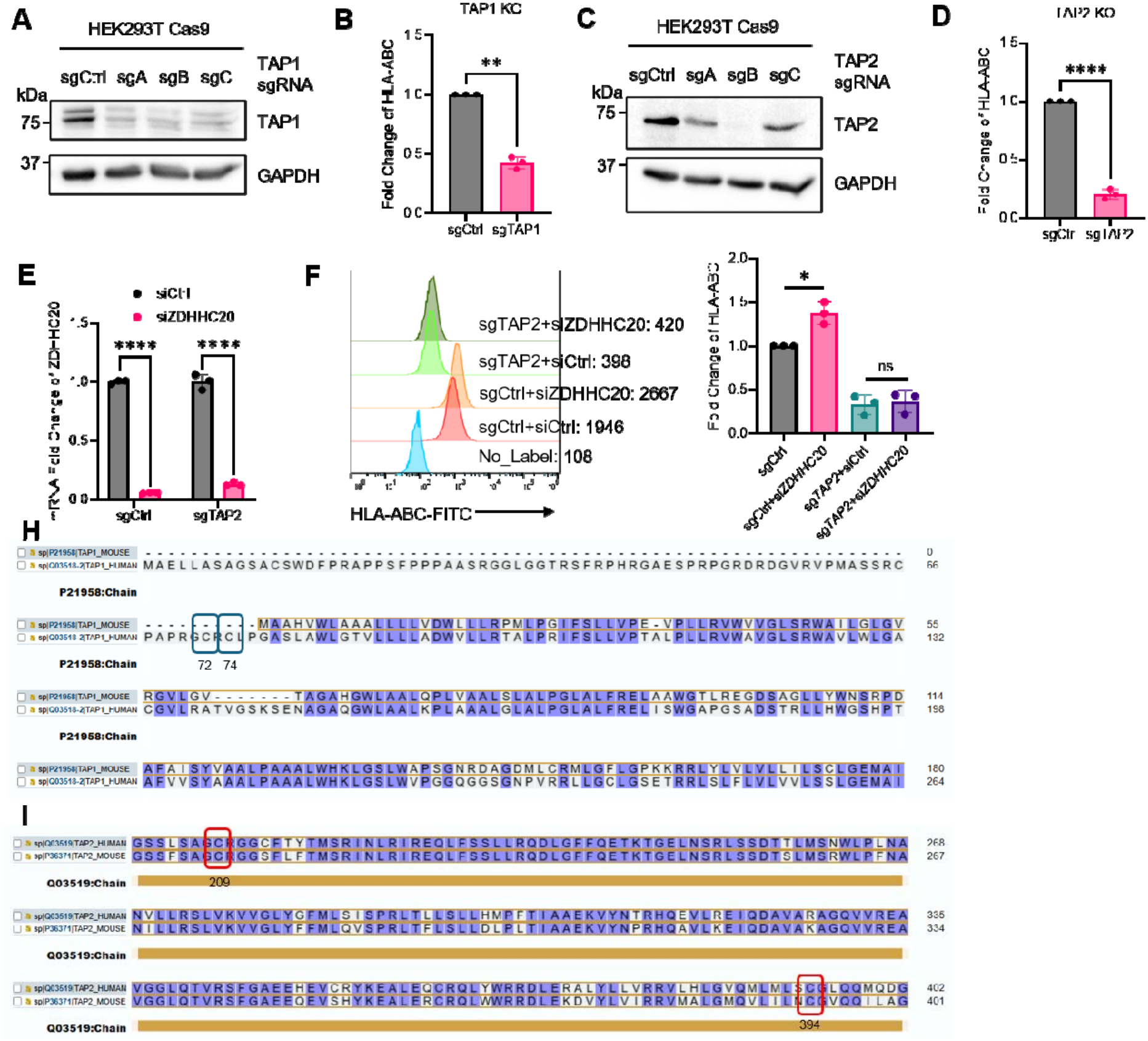
**A**. Generation of TAP1 KO in HEK293T cells. sgA clone is used in other experiments. **B**. Flow-cytometric analysis of surface MHC-I levels in TAP1 KO cells (N = 3). **C**. Generation of TAP2 KO in HEK293T cells. sgB clone is used in other experiments. **D**. Flow-cytometric analysis of surface MHC-I levels in TAP2 KO cells (N = 3). **E**. qPCR validation showing *ZDHHC20* was transiently KD in TAP2 KO HEK 293T cells. **F**. Representative images and quantification of flow cytometry results of *ZDHHC20* transiently KD effect in *TAP2* KO or Control HEK293T cells. **G-H.** Sequence alignment of human and mouse TAP1 (**G**) and TAP2 (**H**). Cysteines 72/74 in TAP1, which are not conserved in the mouse orthologs, are highlighted with blue boxes; cysteine 209/394 in TAP2, which are conserved, are highlighted with red boxes. Statistical analysis: paired t test: (B, D, F); unpaired t test: (E). Sample sizes: (B, D, E, F) N=3.

**Figure S7.**
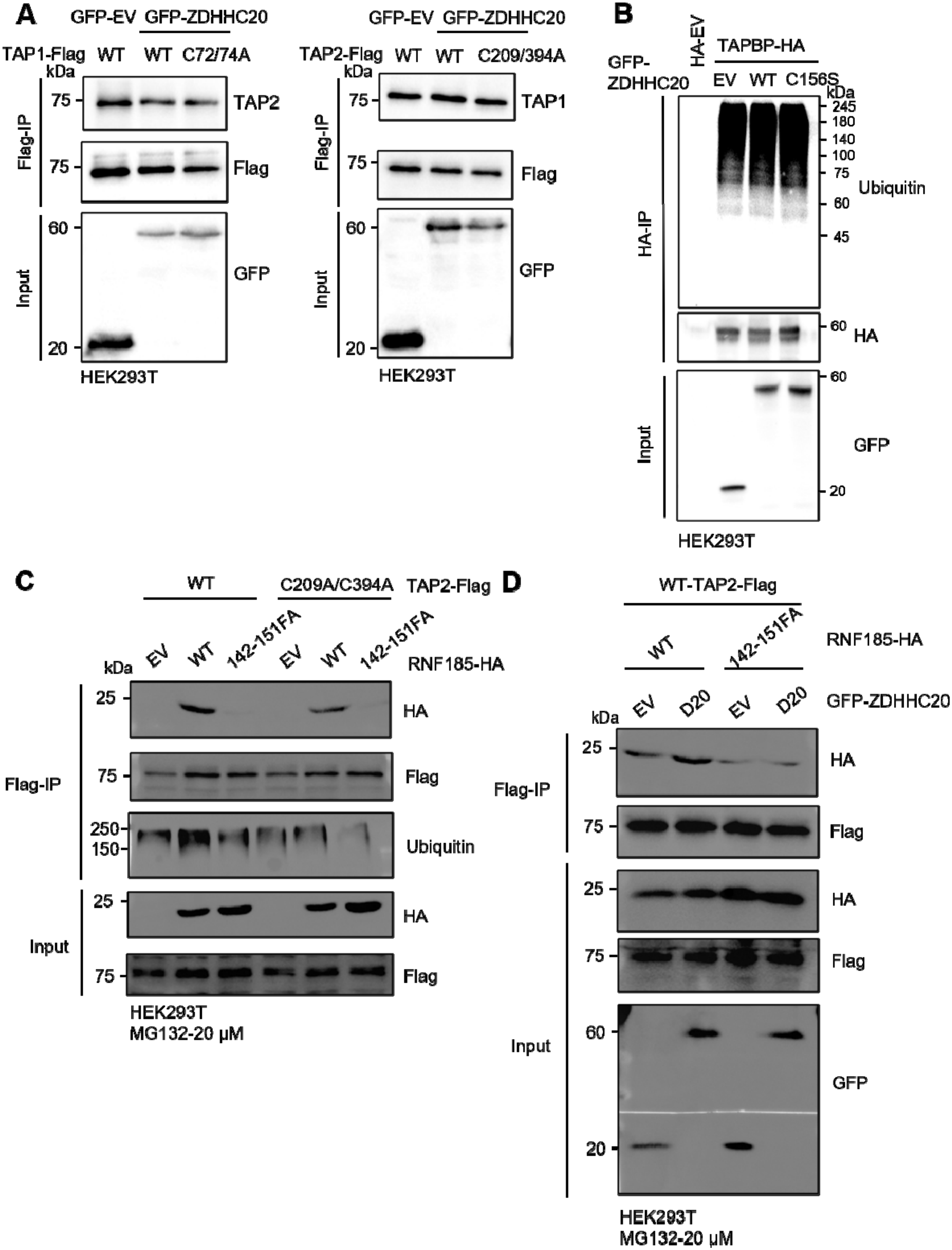
**A.** Co-IP experiments for TAP1/2 complex interaction using WT and palmitoylation deficient mutants of TAP1 and TAP2. Overexpression of ZDHHC20 or mutation of the palmitoylation sites did not affect TAP1–TAP2 complex formation. Input lysates were immunoblotted for GFP-ZDHHC20 expression. **B.** Ubiquitination assay of TAPBP in HEK293T cells expressing HA-tagged TAPBP together with GFP-EV, GFP-ZDHHC20-WT, or the catalytic inactive mutant GFP-ZDHHC20-C156S. Ubiquitination level of TAPBP did not change with overexpression of ZDHHC20. **C**. Co-IP analysis of the interaction between TAP2 and RNF185. HEK293T cells were co-transfected with Flag-tagged WT or palmitoylation-deficient TAP2 (C209A/C394A) together with HA-tagged WT RNF185 or the substrate recognition mutant RNF185-142–151FA. D. Co-IP analysis of the interaction between TAP2 and RNF185-WT or 142–151-FA mutant with or without overexpression of ZDHHC20. Overexpression of ZDHHC20 increased the interaction between TAP2 and WT RNF185 while not the 142_151_FA mutant of RNF185. Image in A, B, C, D shows representative results from two independent experiments.

**Figure S8.**
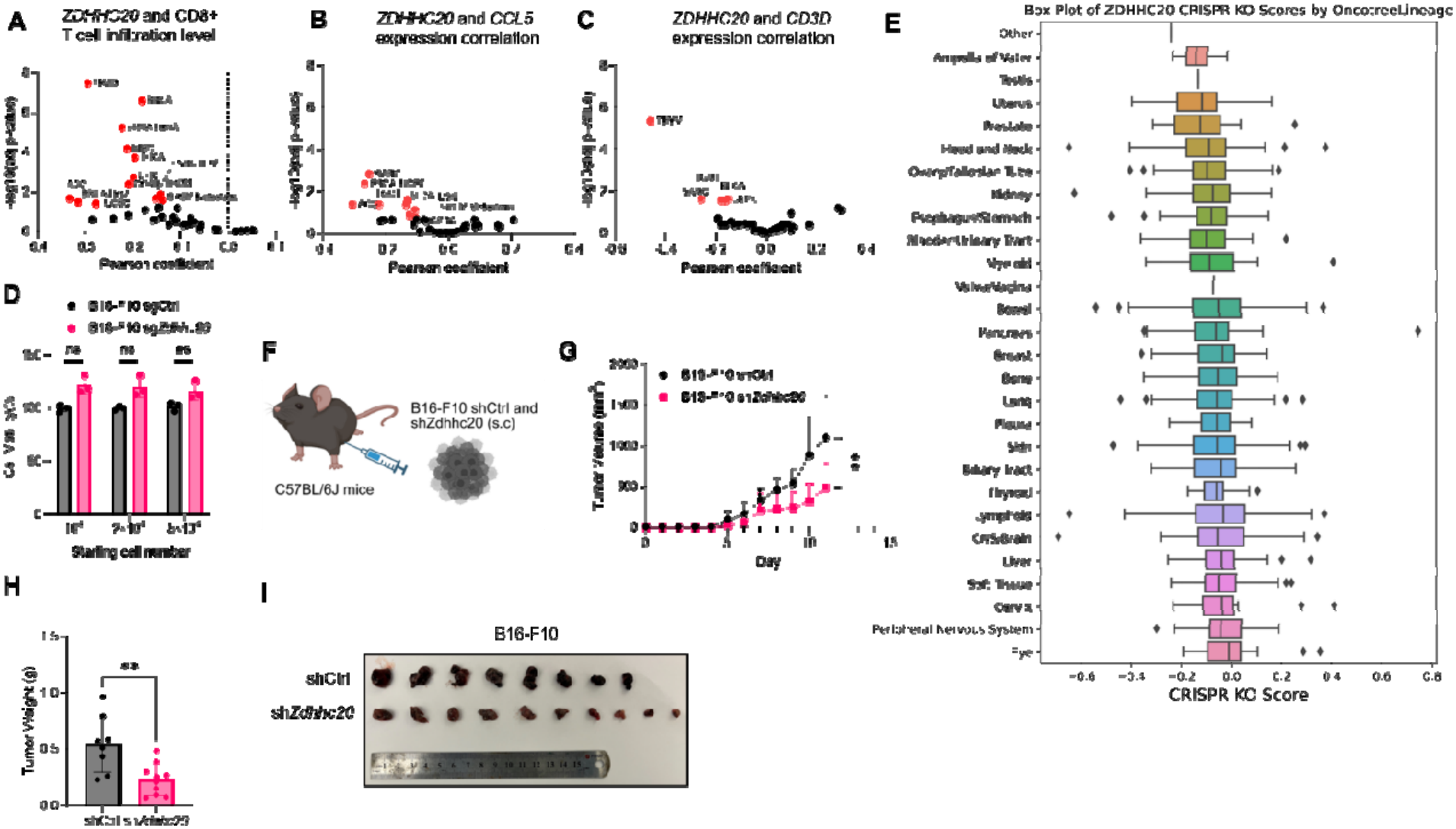
Correlation analysis of tumor *ZDHHC20* levels and CD8^+^ T cell infiltration levels in different cancer types. **A-C**. Correlation analysis of tumor *ZDHHC20* level and CD8^+^ T cell infiltration (**A**)/*CCL5*(**B**)/*CD3D*(**C**) level in different cancer types. **D**. In vitro growth rates of sgCtrl and sg*Zdhhc20* B16-F10 cells. **E**. Depmap analysis of *ZDHHC20* CRISPR KO score in different OncotreeLineages. **F**. Subcutaneous tumor model using shCtrl and sh*Zdhhc20* B16-F10 cells in C57BL/6J mice (N = 8 for shCtrl, N = 10 for sh*Zdhhc20*). **G**. Tumor growth curve. **H**. Tumor weight. **I**. Tumor images of sgCtrl and sh*Zdhhc20* B16-F10 subcutaneous tumors are shown. Statistical analysis: Unpaired t test: (D, G, H).

**Figure S9.**
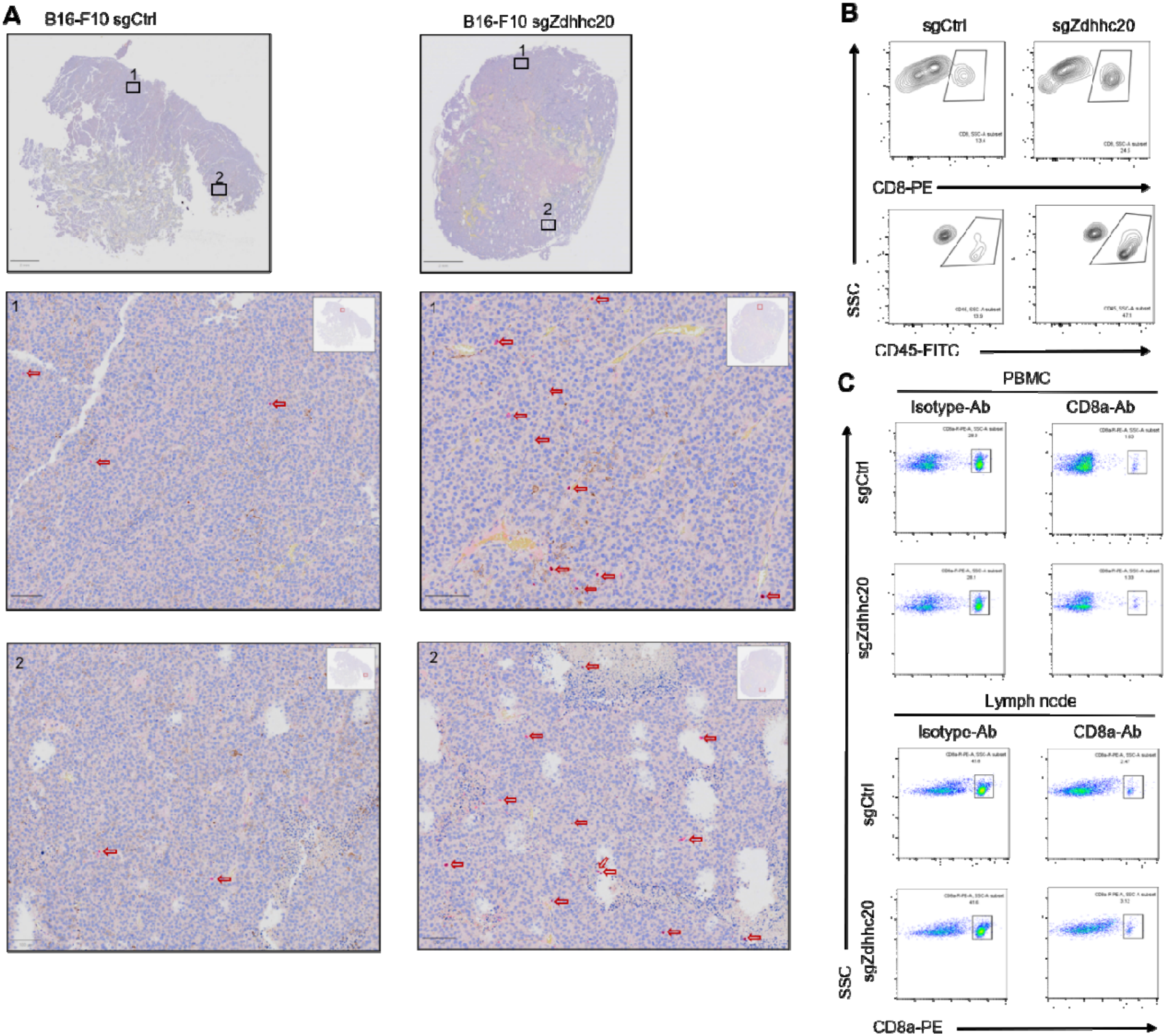
**A**. Immunohistology analysis of tumor infiltrating CD8^+^ T cells in sgCtrl and sg*Zdhhc20* B16-F10 tumor samples. CD8^+^ T cells are labeled with red arrows. **B**. Representative flow cytometry image of tumor infiltrating CD8^+^ and CD45^+^ cells. **C**. Representative flow cytometry images of CD8^+^ T cells in PBMC and lymph nodes after anti–CD8a-Ab treatment.

**Figure S10.**
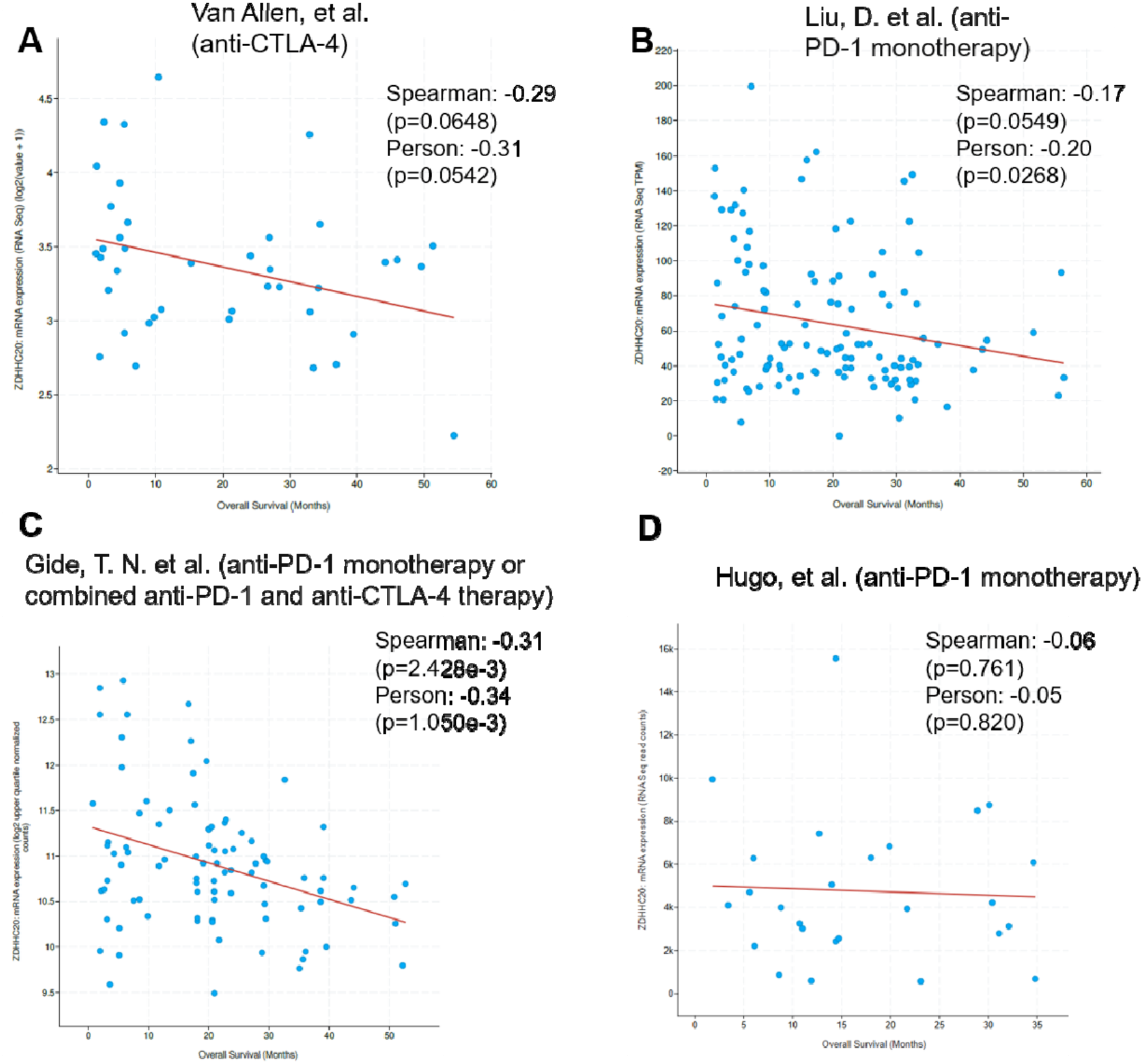
Correlation between *ZDHHC20* mRNA expression and overall survival acros independent anti-PD-1/anti-CTLA-4-treated melanoma cohorts. Scatter plots showing th relationship between *ZDHHC20* mRNA expression and overall survival (months) in four independent, publicly available melanoma cohorts treated with immune checkpoint blockade. (A) Van Allen et al. cohort (anti-CTLA-4; n = 40). (B) Liu et al. cohort (anti-PD-1 monotherapy; n = 121). (C) Gide et al. cohort (anti-PD-1 monotherapy or combined anti-PD-1/anti-CTLA-4 therapy; n = 73). (D) Hugo et al. cohort (anti-PD-1 monotherapy; n = 28). Each point represents an individual patient; red lines indicate linear regression fit. Spearman’s and Pearson’s correlation coefficients with corresponding p-values are shown for each cohort. A negative correlation between ZDHHC20 expression and overall survival was observed in the Van Allen, Liu, and Gide cohorts, reaching statistical significance in the Liu (Pearson p = 0.0268) and Gide (Spearman p = 2.428×10 ³; Pearson p = 1.050×10 ³) cohorts, but not in the Hugo cohort (p > 0.7 for both measures).

**Figure S11.**
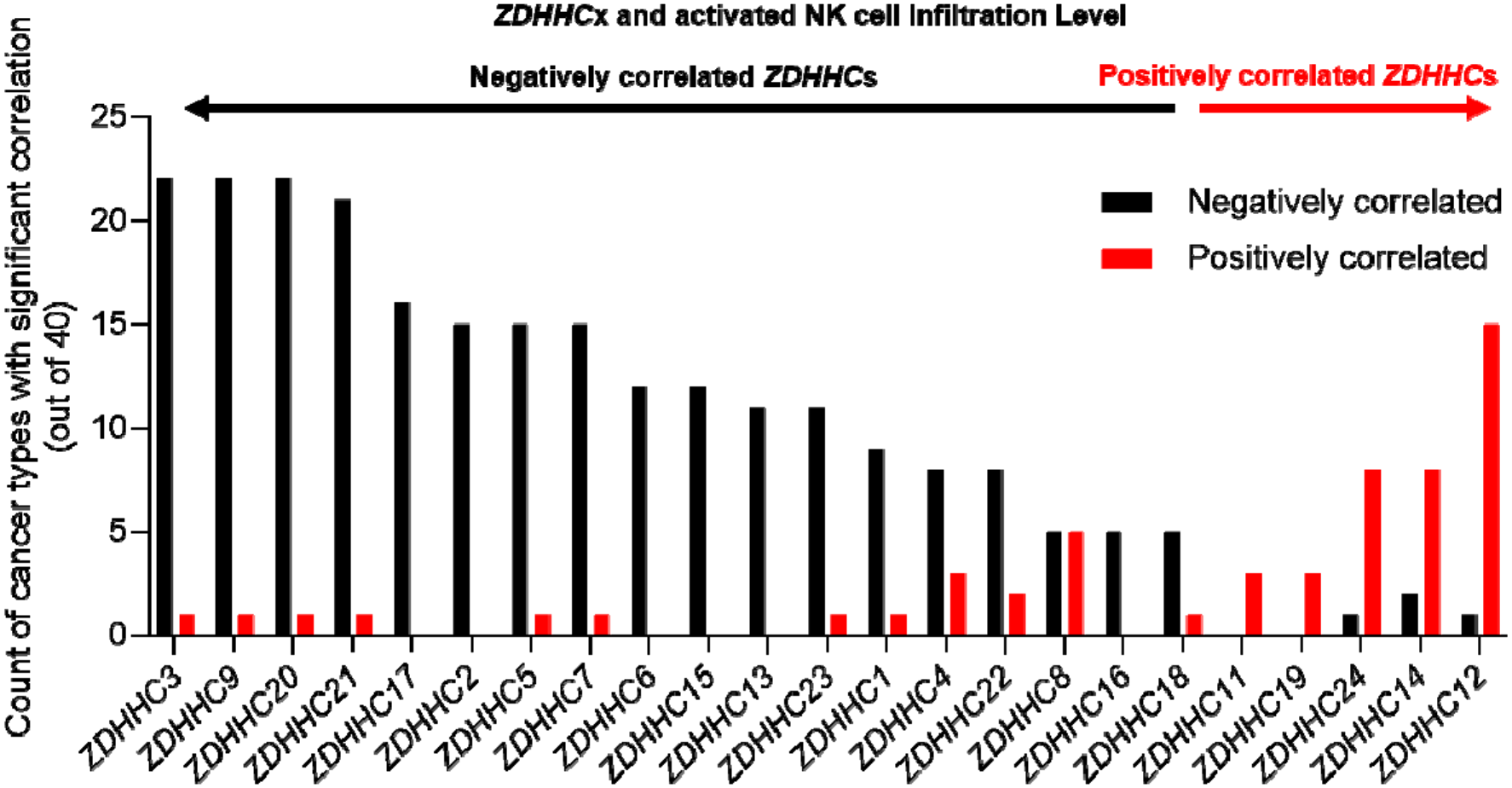
Correlation analysis between tumor infiltrating activated NK cells and expression of each ZDHHC family gene across cancer types. The number of cancer types showing a negative correlation is plotted for each *ZDHHC* gene.

**Figure S12.**
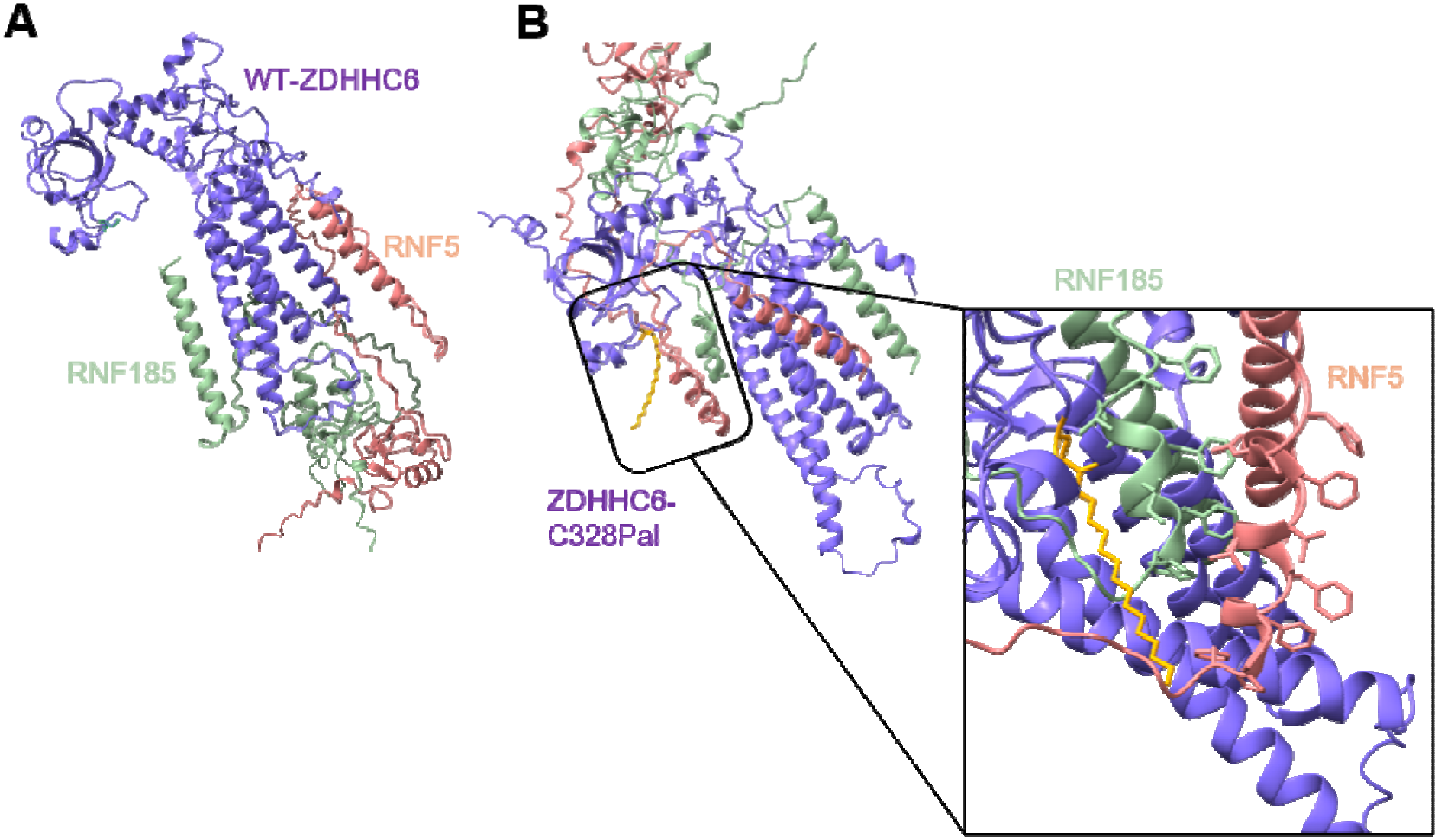
AlphaFold 3 prediction of ZDHHC6 binding pose with RNF5/RNF185. **A**. AlphaFold3-predicted structure of WT ZDHHC6 (purple) in complex with RNF5 (salmon) and RNF185 (green). **B**. AlphaFold3 model of C328-palmitoylated ZDHHC6 in complex with RNF5 and RNF185. The inset highlights the proximity of the phenylalanine-rich domains of RNF5 and RNF185 to the palmitoylated Cys328 residue (yellow), suggesting that palmitoylation at Cys328 may facilitate the interaction of ZDHHC6 with these E3 ligases.

